# Assessing the potential of bee-collected pollen sequence data to train machine learning models for geolocation of sample origin

**DOI:** 10.64898/2026.03.29.715128

**Authors:** Rebecca A. Hayes, Andrew D. Kern, Lauren C. Ponisio

## Abstract

Pollen is a robust and widespread substance that captures a historical snapshot of a specific time and place, and it can be used to track movements through space by examining the pollen deposited on various objects. Palynology, the study of pollen, is used across fields such as conservation, natural history, and forensics, where it is particularly useful for tracing the origin and movement of objects. However, pollen has remained underutilized due to the difficulty of distinguishing many pollen taxa beyond the family level and limited pollen reference material to support location predictions. With recent developments in pollen DNA metabarcoding these issues have been rectified, but much of the available pollen data are primarily from wind-pollinated species, which are widespread and less informative of specific sample locations. Bee-collected pollen presents an untapped resource in training predictive models to geolocate sample origin. Here we compiled bee-collected pollen DNA sequence relative abundance data from three projects in the western U.S. and assessed the accuracy of supervised machine learning models to predict the location of sample origin based solely on pollen assemblage, without the need of incorporating additional data. Random Forest and k-Nearest Neighbors models yielded high accuracy across all projects. We also found that models trained on taxonomically clustered pollen assigned sequence variants (ASVs) performed slightly better than those trained on raw sequence data, but the difference was minor, indicating that models trained on raw sequence data can reliably predict location and avoid the time-consuming taxonomic assignment process. Our results demonstrate the utility of repurposing bee-collected pollen for geolocation and provide a framework for employing supervised machine learning in future geolocation efforts.

**Highlights:**

- Bee-collected pollen metabarcoding data was used to accurately predict sample origin
- Random Forest and k-Nearest Neighbors algorithms were most accurate with lowest error
- Taxonomically-classified and raw DNA sequence data training sets performed comparably

## 1 Introduction

Pollen stands out among other biological materials for its durability and ubiquity across environments and seasons, which allows investigators to peer back in time to a particular spatiotemporal context by examining pollen taxonomy, dispersal, and its relative abundance in time and space. Palynology, the study of pollen grains and composition, has been successfully used to understand paleobotanical communities, past climates, archaeological artifacts, historical movement of populations and objects, and to inform forensic investigations (Davis, 1969; Bryant Jr and Holloway, 1983; Faegri and Iversen, 1992; Mildenhall et al., 2006). It is particularly applicable to situations involving suspected transfers of objects between locations or to those that occurred in areas with distinctive plant communities (Bell et al., 2016a). However, despite its demonstrated strong potential as a tool to understand history and movement, palynology has not been widely adopted by researchers across fields that would benefit from the information pollen provides. Historically, palynology required collaboration with experts in pollen identification by morphology, which posed a barrier to adoption due to the scarcity of trained researchers (Wiltshire, 2016; Bryant and Jones, 2006). Moreover, expertise is often geographically restricted to the plant communities in which an individual was trained and is not immediately generalizable to other plant communities (Wiltshire, 2016). Traditionally, palynological reference libraries have been built by investigators for a specific use-case and apply only to a very particular situation, thus are not readily reused for other cases (Wiltshire, 2016). Image-based morphological reference libraries have been developed (e.g. PalDat Palynological Database; Weber and Ulrich, 2017), however such libraries require individual researchers to upload their pollen images to the database and are biased towards well-studied plants and systems. Additionally, the pollen of many plants is only visually distinguishable to the family or subfamily level: for example, the widespread families Fabaceae, Lamiaceae, and Apiaceae are difficult to identify beyond the family level (Erdtman, 2023). Thus, utilizing such resources is time consuming and may not be resolvable to a useful level. Due to these roadblocks, the future of palynology would benefit from the use of modern techniques that are more widely taught across disciplines and can provide deeper taxonomic resolution.

DNA sequencing-based approaches represent a revolutionary advancement for many fields that examine species composition using molecular techniques, enabling rapid and robust analysis of biological evidence (Yang et al., 2014). In the context of palynology, improvements in molecular tools, such as DNA metabarcoding, have facilitated the study of plant communities over time and space without relying on morphological characteristics (Keller et al., 2015). For example, primers that amplify highly conserved regions of the plant genome, such as the sequence encoding the large subunit of the photosynthetic enzyme RuBisCo, allow the characterization of multiple pollen species from a single sample (Bell et al., 2017). The molecular skills and equipment required for DNA metabarcoding are more widely taught and readily available in typical molecular laboratories (Bell et al., 2016a). Additionally, plants worldwide can be distinguished using the same primers, transcending the geographic limitations of morphological identification Bell et al. (2016a). Finally, pollen DNA sequence reference libraries, such as those compiled by NCBI and RDP, continuously grow each year as researchers outside the field of palynology deposit sequences from their studies, facilitating taxonomic identification to the genus or species level (Omonhinmin and Onuselogu, 2022). While the methodological capabilities for using DNA metabarcoding to identify pollen have existed for some time, applying these techniques has yet to be fully realized.

To be successfully deployed as a tool in palynology, pollen metabarcoding data must be matched to a specific location based on known local pollen assemblage. DNA sequence reference libraries, such as those provided by NCBI and RDP, expand each year as more reference sequences are deposited; however, samples are not always georeferenced beyond the country level, as this is not a requirement for inclusion in repositories. Previously, paleoecologists have achieved high predictive accuracy using the modern analog technique with cross-validation to produce environmental reconstructions, however this is most successful at the continental- to sub-continental-scale or for predicting vegetation zones of samples (Williams and Shuman, 2008). While distinguishing between continents, countries, or vegetation zones of origin can be useful in certain large-scale cases, geolocation at a more granular scale would expand its potential applications to regional or local investigations. However, until repositories require more specific location information for sequence data, alternative datasets that include pollen sequence data paired with precise geographic location could provide a solution.

Pollinator community ecologists worldwide commonly use DNA metabarcoding of the rbcL gene to study bee diets, generating datasets that can be repurposed for palynological reference libraries (Bell et al., 2016b). In pollen metabarcoding studies, it is standard practice to meticulously record the location information of the specimens from which pollen is collected. This means that pollen DNA metabarcoding data contains all the necessary information to be useful for geolocation applications. Beyond its location data, bee-collected pollen offers additional benefits for geolocation applications. Paleoecologists have extensively studied sediment pollen assemblages, however sediment pollen is typically dominated by wind-dispersed pollen types, whereas entomophilous pollen is underrepresented (McGlone, 1988). Because wind-dispersed pollen travels large distances, is widespread spatially, and is produced in large quantities (Di-Giovanni and Kevan, 1991), it may be less informative for forensic studies seeking to pinpoint a particular location. Thus, geolocation can benefit from pollen assemblages that are highly temporally and spatially specific, such as bee-collected pollen. Bees forage in many different environments, both natural and human-modified, spanning from remote to rural to urban, and have temporal variation in foraging behaviors between individuals (Lemanski et al., 2019). For example, the western honey bee (*Apis mellifera*) is the most widespread introduced pollinator, spanning across many global agricultural hotspots and human population centers (Visick and Ratnieks, 2023). Social bees, such as the generalist honey bee, exhibit foraging patterns that capture a representative snapshot of surrounding flowering plant communities (Young et al., 2021). Additionally, the foraging patterns of individuals scale with plant spacing, also providing good spatial representation of a local plant community (Morris, 1993). Similar to honey bees, bumblebee species (*Bombus* spp.) as individuals tend towards fidelity for a few plant species, but at the colony level, each individual typically varies in preferred forage (Heinrich, 1976). Incorporating samples from a wide variety of bee species, which vary widely in floral diets, is also common in pollen metabarcoding studies, providing increased coverage of the local plant diversity to better inform models (Cappellari et al., 2022). As funding agencies increasingly adopt open science practices, these datasets are increasingly made available to the public for download (Elger et al., 2016). The outstanding challenge, then, is to determine how best to model the relationship between pollen assemblage and location.

Even with precise and accurate geolocation data attached to pollen assemblage data, modeling their spatial relationship is not a trivial task. We lack a clear and comprehensive understanding of the distribution of bee-collected pollen communities and the mechanisms of pollen deposition, which complicates modeling. Several strategies have shown promise in rectifying both issues, including the incorporation of species distribution modeling, historical sample data from museum specimens, spatial optimization approaches, and network-based approaches as a precursor to more complex models to better link pollen to place (Goodman et al., 2015; Warny et al., 2020; Tong et al., 2021; Helderop et al., 2021). While each of these techniques improves upon the original methods based on morphological identification, none provides a stand-alone, simple workflow that incorporates pollen DNA sequence data and can be quickly adopted by practitioners who may not have a strong statistical background or access to historical datasets.

For optimal usefulness in palynology, an algorithm that predicts geographic location from pollen sequence data is ideal. However, many unknown parameters influence which pollen species are likely to be present in a mixed sample, making it difficult to build a model for this purpose. Supervised machine learning, which relies solely on training data without the requirement of additional inputs and can handle spatially-heterogenous data, can bridge this divide (Du et al., 2020). These methods are increasingly used in ecology to untangle complex relationships between ecological communities (Thessen, 2016). Moreover, machine learning algorithms have proven useful in a wide variety of geolocation tasks including recovering location from text-based event data, determining collection location from microbial metabolic fingerprinting, predicting location of origin from genetic variation, and informing urban planning by using social media data to predict vehicle crashes (e.g. Lee et al., 2019; Huang et al., 2020; Battey et al., 2020; Milusheva et al., 2021). In palynology, machine learning has improved classical morphology-based pollen identification by combining computer vision with plant phenological data, thereby streamlining pollen classification and increasing the geographic resolution of predictions (Hwang et al., 2013). However, training computer vision models still requires accurate morphological classification of representative pollen images, necessitating the involvement of expert pollen microscopists. Employing supervised machine learning algorithms that make predictions based on training datasets organized in data frames rather than raster images can circumvent the need for morphological classification altogether. Such models have been employed in a palynological context to predict plant distributions using species occurrence data and climatic variables, with the ultimate goal of using network analysis to narrow down the possible origin locations for a sample (Helderop et al., 2021). However, that approach was intended as a first-pass filter to help reduce the computational intensity required by more complicated image-based predictive algorithms. Building predictive workflows for geolocation presents significant challenges, primarily due to the computational demands and time required for complex algorithms to model spatial relationships within the data. Machine learning techniques often outperform traditional statistical modeling in both speed and accuracy. For instance, the Locator deep learning method, which predicts location based on individual genotypic data, achieves highly accurate geographic predictions in minutes, compared to hours with the leading statistical modeling approach (Battey et al., 2020). Locator was trained on a genotype information per sample; however, for palynological applications, predictions would ideally be made from a collection of genotypes, i.e., the composition of pollen metabarcoding data. In the case of Locator, the genome of each sampled individual reflects the geographic spread of their ancestors, providing sufficient variation to place the individual at a particular location (Battey et al., 2020). Though individual metabarcoded sequences are a small portion of the whole genome and therefore contain less information in an individual sequence than deeper sequencing techniques, the particular composition of different pollen species contained in a single sample provides information on local plant communities that vary spatially. To our knowledge, no studies have examined the utility of supervised machine learning models in geolocating samples using community pollen DNA sequence data.

Here, we explore whether the location of the sample origin can be accurately predicted from mixed-species pollen DNA sequence data with machine learning. We train models using bee-collected pollen from three discrete projects across the western United States as a test case to demonstrate the value in repurposing the copious amounts of available pollen DNA sequence data amassed from pollinator studies. We compare the strengths and weaknesses of using classified and raw pollen sequence data to train six popular machine learning algorithms, providing a framework for future DNA sequence-based palynological applications. This work provides a pathway for employing machine learning techniques and DNA sequence data for palynology, a critical step toward expanding access and enhancing efficiency in the subfield.

## 2 Methods

### 2.1 Sample collection and study sites

To train our geolocation models, we compiled a training dataset of pollen from bees collected across three projects in California, Arizona, New Mexico, and Oregon, representing a range of environmental types, geographic locations, and floral communities. We compiled the training data across three datasets to increase model generalization to a larger geographic extent and because practitioners compiling training data for geolocation from publicly-available repositories would likely draw from multiple projects to better refine geographic predictions. Additionally, to emphasize the utility of bee-collected pollen reference libraries, we selected projects that encompass the range of typical studies that generate georeferenced pollen sequence data from bees.

#### 2.1.1 Sky Islands Project

The team collected samples during surveys of 7 isolated, high-elevation meadows in 2018 and 2021 in the Madrean/Cordilleran Sky Islands in Arizona and New Mexico (Smith et al., 2021). Sites were relatively unmodified by human activity and represent natural, highly-diverse floral communities. Individual sites along a floral biodiversity gradient showed greater floral diversity at the southernmost sites than at the northernmost sites. The most abundant bee species we collected for this project were *Apis mellifera*, *Bombus huntii*, and *Bombus centralis*. The closest meadows were separated by 1.52 km and the farthest by 553.5 km. The spatial extent of the sites sampled, estimated as the area of the minimum convex polygon enclosing all sampling locations for this project, was approximately 52276 *km*^2^. In total, this dataset included 228 pollen samples.

#### 2.1.2 California Sunflower Project

The team collected samples from the California Sunflower project during surveys of mass-flowering sunflower fields in Yolo County, California, in 2019 (Cohen et al., 2021). The landscape was primarily intensively managed row crops, some of which had flowering, unmanaged, weedy field margins and native plant hedgerows. The most abundant bee species we collected for this project were *Melissodes agilis*, *Lasioglossum incompletum*, and *Apis mellifera*. Individual sites were geographically close relative to the other projects. The spatial extent of the sites sampled, estimated as the area of the minimum convex polygon enclosing all sampling locations for this project, was approximately 328 *km*^2^. In total, this dataset included 1178 pollen samples.

#### 2.1.3 Pacific Northwest Forests Project

The team collected samples from the Pacific Northwest Forests project during surveys of bee communities in post-fire and managed timber plantations from 2020-2023 in the Coast and Cascade Mountain Range (Fan Brown et al., 2025). This project included sites in multiple high-severity fire scars that originally burned in 2020 including the California Dixie Fire and Oregon Holiday Farm and Beachie Creek Fires. Within a fire, sites were separated by at least 1 km. Additionally, sites in the Coast Mountain Range in Oregon were unburned but managed for timber production (Fan Brown et al., 2025). Harvested forest sites were separated by at least 1 km. The most abundant bee species we collected for this project were *Bombus vosnesenskii*, *Bombus caliginosus*, and *Bombus sitkensis*. The spatial extent of the sites sampled, estimated as the area of the minimum convex polygon enclosing all sampling locations for this project, was approximately 95907 *km*^2^. In total, this dataset included 176 pollen samples.

The spatial extent of the sites sampled, estimated as the area of the minimum convex polygon enclosing all sampling locations for this project, was approximately 986309 *km*^2^. For samples from all three projects, we collected bees with hand nets and stored them in sterile collection tubes on dry ice until returning to the lab, upon which they were stored in a −80^◦^*C* freezer. For individuals carrying pollen, we collected scopal pollen separately from the specimens into 1.5mL microcentrifuge tubes using sterile technique and stored them dry at −80^◦^*C* until DNA extractions. We also completed vegetation surveys for all three projects to document the plant diversity at each site to determine feasible pollen genus and species identifications.

### 2.2 Pollen DNA extraction and library preparation

We used Machery-Nagel NucleoSpin 96 Food kits (Dü ren, Germany) to extract pollen DNA from the samples, following the provided kit protocol. To each sample tube, we added 550 *µ*L of warmed buffer CF, 10 *µ*L of Proteinase K, and approximately 100 *µ*L of 0.1 mm zirconia beads before lysis using a Qiagen Tissue Lyser II. Samples were transferred to 96 well plates after the first centrifuge spin step, then we followed the remaining steps from the kit protocol. We included a negative control well in each plate to avoid contamination.

We used the Illumina MiSeq platform to identify extracted pollen DNA (2*x*300, *V*3 reagents) (Illumina, Inc, San Diego, CA) following a dual-index, inline barcoding approach (McFrederick and Rehan, 2016). We amplified 180-220 bases of the plant rbcL gene using the RBCL7 CTCCTGAMTAYGAAACCAAAGA, and the RBCL8 GTAGCAGCGCCCTTTGTAAC primers McFrederick and Rehan (2016). The rbcL gene codes for a portion of the enzyme RuBisCo, a necessary enzyme in photosynthesis common to all land plants (1 et al., 2009). Despite the highly conserved nature of the rbcL gene, variation between taxa has been successfully used to distinguish plants in mixed pollen samples using the same primers and NGS (Bell et al., 2019).

We completed Illumina library preparation in four steps: a PCR round to produce rbcL amplicons, an enzymatic cleaning step, a second PCR round to add Illumina adapters, and, finally, a normalization step to standardize across sample wells and plates. The first-round PCR reactions used 4 *µ*L of DNA, 19 *µ*L ultrapure water, 10 *µ*L Phusion High-Fidelity Master Mix (Thermo Fisher Scientific, Waltham, MA), and 1 *µ*L each of 5 *mu*M primer stock. We used an annealing temperature of 52^◦^C for 35 cycles. Negative controls were included in each reaction to detect potential contaminants. Enzymatic PCR cleanup was performed using ExoSAP-IT PCR Product Cleanup Reagent (ThermoFisher Scientific, Waltham, MA) to remove excess primers and unincorporated dNTPs using the following protocol: we incubated samples at 37^◦^C with 0.025 *µ*L Exonuclease I, 0.25 *µ*L Shrimp Alkaline Phosphatase, and 9.725 *µ*L ultrapure water before inactivating the enzyme at 95^◦^C for 5 min. To add Illumina adapters to our cleaned amplicons in a second PCR step, we used HPLC purified PCR2F and PCR2R primers, as in Kembel et al. (2014): CAAGCAGAAGACGGCATACGAGATCGGTCTCGGCATTCCTGC and AATGATACGGCGACCACCGAGATCTACACTCTTTCCCTACACGACG. We annealed the samples at 58^◦^C for the second round of PCR reactions, with 35 cycles. We normalized amplicons using 18 *µ*L of PCR product in SequalPrep Normalization plates (ThermoFisher Scientific, Waltham, MA). We pooled five *µ*L from each normalized sample for sequencing. Libraries for each of the three projects were sequenced with the MiSeq Reagent Kit v3 with 2 × 300 cycles in two distinct runs for the Sky Islands project, five distinct runs for the Sunflower Monoculture project, and two distinct runs for the PNW Forests project.

### 2.3 Pollen bioinformatics

We used QIIME 1 (2019.1) to parse paired-end in-line barcodes, then used QIIME 2 (2019.1) to demultiplex reads and filter rbcL sequence libraries (Bolyen et al., 2019). First, we visualized and trimmed the low-quality ends of the reads with QIIME2, then merged data from multiple Illumina runs across projects. We used DADA2 (Callahan et al., 2016) to remove chimeras, remove reads with more than two expected errors, and assign sequences into amplicon sequence variants (ASVs).

We classified pollen ASVs in two ways following bioinformatic protocols in Smith et al. (2024): with a pre-trained rbcL RDP classifier (Bell et al., 2017) and with NCBI BLAST (code available at https://github.com/hayesrebecca/pollenGeolocation). For the RDP classifier, we followed the protocols in Bell et al. (Bell et al., 2017). To classify using BLAST, we first built a local rbcL database by downloading all rbcL nucleotide sequences from NCBI and compiling it into a database using BioPerl (Stajich et al., 2002). We used a custom BioPerl script to run local BLASTn searches and to parse the results, keeping the top BLAST hit as the call for NCBI (Smith et al., 2024). We assigned the final taxonomic designations manually by comparing the output of these two classifiers. Where RDP and NCBI classifiers agreed on genus and species, we assigned that species. If RDP and NCBI agreed only on the genus but not on the species, we accepted the genus. We examined our vegetation survey data to determine whether both, one, or neither species was present at the sites. If only one species was present, we assigned it to that species. If both species were present, we chose the NCBI classification. If neither species was present, we examined the USDA Plants Database for the region to determine whether that species is feasible for that location, and, if so, assigned it. If that species was not listed in the USDA Plants Database for that region, the final call was made to the genus level. If RDP and NCBI classifiers disagreed at both genus and species levels, the final call was made at the lowest taxonomic level, with consensus between RDP and NCBI.

We used the MAFFT aligner (Katoh and Standley, 2013) and FastTree v2.1.3 in QIIME 2 to generate a phylogenetic tree of our sequences (Price et al., 2010). We filtered the resulting ASV table for features corresponding to contaminants identified in the blanks (Karstens et al., 2019) or present in only one read. To account for variable sequencing depth, we generated an alpha rarefaction curve for each sequencing run and subsampled.

We merged the relative abundance ASV tables from the three projects into a unified data frame for analysis. Before the merge, the Sky Islands project included 25 unique pollen taxa, the California Sunflower project included 120, and the Pacific Northwest Forests project included 69. This resulted in a training dataset of relative abundances for 185 pollen taxa across all projects due to some overlap in clustering between projects (Fig. S 1).

Additionally, to understand how taxonomic assignment can influence predictions, we generated a second unified data frame that retained the raw DNA sequences for each ASV rather than the assigned taxonomy. This represents the most conservative use case, in which closely related sequences were not clustered and instead treated as unique strains of pollen. The Sky Islands project included 240 raw pollen sequences, the California Sunflower project included 566 raw pollen sequences, and the Pacific Northwest Forests project included 189 raw pollen sequences. This resulted in a training dataset of relative abundances for 954 raw pollen sequences across all projects due to some shared sequences among projects, particularly between the California Sunflower project and some of the Californian sites in the Pacific Northwest Forests project (Fig. S 2). We trained all machine learning models separately on both taxonomically assigned data and raw sequence data for comparison.

For both raw and taxonomically clustered ASVs, it was more common for ASVs to be restricted to a few sites rather than widespread (Figs. S 3 and 4). However, across all three projects, ASVs were detected at all sites (Figs. S 3 and 4).

### 2.4 Selecting and training machine learning models

To understand how well geographic location can be estimated from the DNA composition of pollen samples, we evaluated a suite of common machine-learning algorithms. These were chosen based on demonstrated success in modeling multi-output relationships in tabular datasets (Thessen, 2016). Supervised learning base models included MultiTaskLasso, Support Vector Regression, k-Nearest Neighbors, and decision trees, all as implemented in scikit-learn (Pedregosa et al., 2011). Lasso regression is a type of linear regression that includes a penalty term on the regression equation to regularize and prevent overfitting (Ranstam and Cook, 2018). Support vector machines work in a high-dimensional space to find the hyperplane that best separates continuous target values (Awad et al., 2015) with extensions to non-linear regression. k-Nearest neighbors regression uses proximity to neighboring points within a fixed radius to predict the identity of target variables (Steinbach and Tan, 2009). Decision tree regression uses binary splits of input features, making stepwise decisions to predict target variables and minimize Gini impurity (Testas, 2023).

We also tested the performance of several ensemble algorithms, which integrate a collection of weak learners to produce an ‘ensemble’ prediction, using randomly selected feature subsets for each ensemble member (Dietterich, 2000). Ensemble learning algorithms often yield improved accuracy and generalization to new data, as well as reduced overfitting and bias (Dietterich, 2000). These included random forests and XGBoost. Random forest regression utilizes multiple decision trees in parallel and combines their predictions to determine the best decision path for predicting targets (Breiman, 2001). XGBoost, or extreme gradient boosting, similarly uses multiple decision trees, but rather than generating trees in parallel, it sequentially updates subsequent trees using the previous tree to inform the best decision path (Chen et al., 2015).

We conducted all downstream data cleaning and analysis in Python (version 3.11.14). We split the merged relative abundance ASV table into features representing each ASV and targets representing the latitude and longitude coordinates of each sample. We divided the dataset into training and test sets using the train test split function from the scikit-learn module, reserving 20% of the data from each project for the test dataset. We conducted the train–test split separately for each project to ensure that the test dataset preserves the relative balance of data between projects. We standardized features and targets separately for training using the StandardScaler function from scikit-learn, with training and testing data scaled independently to prevent information leakage.

### 2.5 Hyperparameter tuning

To optimize the performance of individual machine learning models, we conducted a randomized grid search using cross-validation across model hyperparameters (Bergstra and Bengio, 2012). While the goal of machine learning is to learn parameter values from the data, many algorithms have specific parameters that describe their complexity and learning speed, which are typically set before training and not learned from the data itself (Rimal et al., 2024). Thus, randomized grid search in combination with cross-validation allows hyperparameter optimization for a particular task and dataset and again insures against leakage. Each model has its own specific number and type of hyperparameters, so we selected values to allow variation in model complexity while avoiding excessive computation from searching an unnecessarily large grid (Table S 1). We performed a randomized grid search with 5-fold cross-validation, repeated 3 times, yielding 15 iterations per model. We randomly selected six distinct hyperparameter combinations for each model, focusing on parameters that control model complexity. In total, the grid search for each model covered 90 unique model fits. We selected the combination of parameter values that minimized RMSE of predicted longitude and latitude and subsequently applied each model using those values to the complete training set (Table S 2).

### 2.6 Model assessment and feature importance

To assess model performance, we calculated the coefficient of determination (*R*^2^), root mean squared error (RMSE), median absolute error (MAE), and average distance loss (AvgDistLoss). For the Random Forest models, we determined feature importance for the top 25 most important features to identify which pollen sequences and ASVs were most influential in predicting latitude and longitude (Qi, 2012).

## 3 Results

Each of the models we tested were able to predict geographic location of sample collection from pollen RBCL sequence data to some degree, whether trained on classified pollen ASVs and raw sequences, however they varied in performance (Table 1, Table 2, Fig. 1). All hyperparameter-tuned models outperformed their untuned counterparts. We found that the taxonomically-clustered pollen ASV training set consistently yielded better predictions than the raw DNA sequence training set. All six of the models trained using the taxonomically-clustered data explained more than 84% of the variation in the held-out data (Table 1, Figure 2), with the k-nearest Neighbors algorithm explaining the highest amount of variation (97.6%) while minimizing RMSE (0.15). The average geodesic error for this model was 10.2 km with a standard deviation of 22.9 km. This predictive error is small relative to the total study area size and each individual project’s extent: the CA Sunflower project, which covered the smallest area by far, included 328.00 *km*^2^. The k-nearest Neighbors algorithm was also the only model that reliably explained any variation in the test dataset on a per-project scale, indicating generalizability across all projects (Table 1, Figure 3). For the models trained on raw pollen DNA sequences, the Random Forest algorithm yielded the best predictions from the held-out data, accounting for 88.2% of the variation in the data (Table 2, Figure 4). The per-project predictive accuracy for the models trained on raw pollen DNA sequences was lower than for models trained on taxonomically-classified data (Table 2, Figure 5). Negative R^2^ values in the per-project evaluations indicate that, although models were trained on the combined dataset, they explain less variance within an individual project than a baseline model predicting that project’s mean, reflecting distributional differences in plant composition.

**Figure 1:**
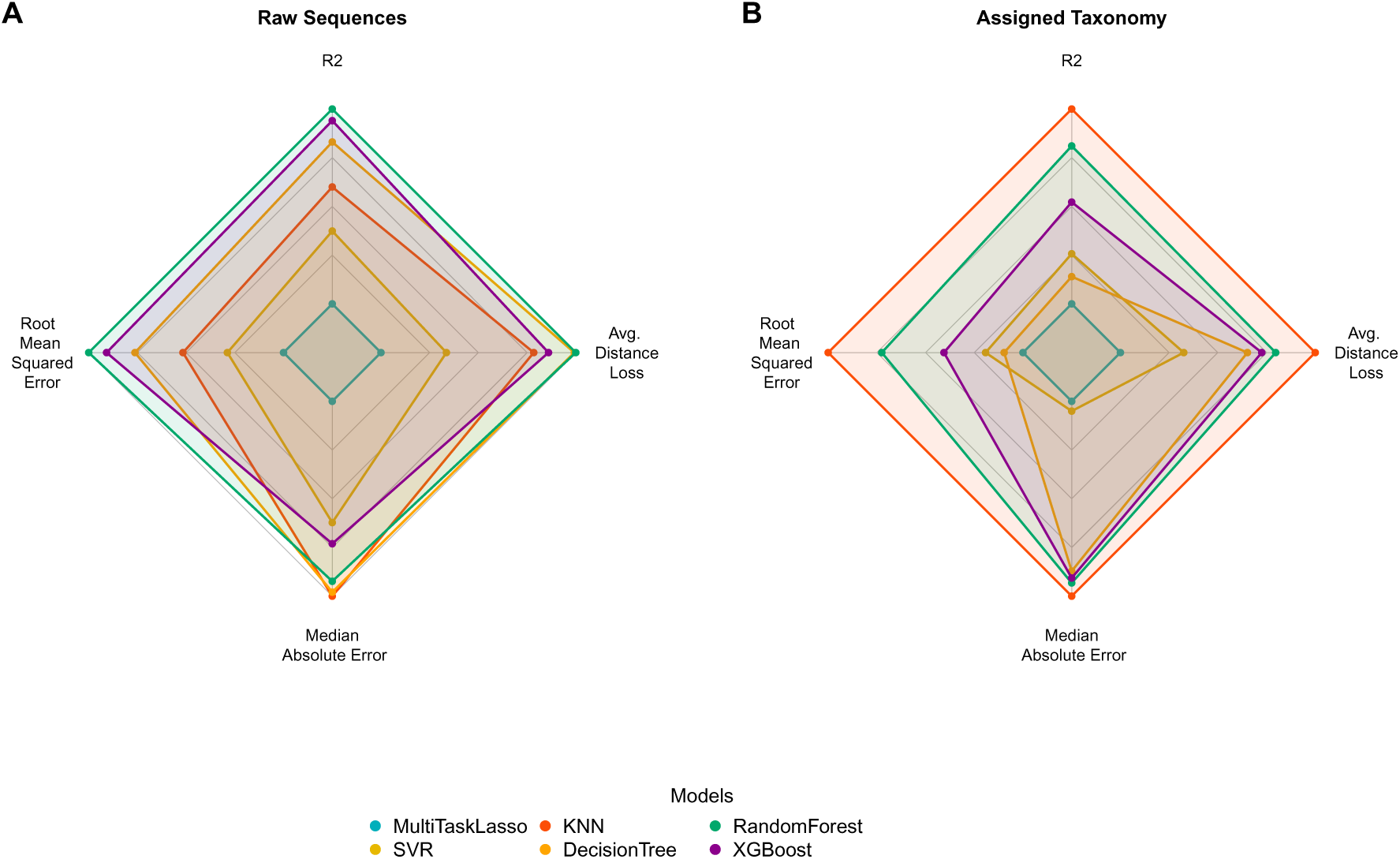
Relative performance between models trained on A) raw pollen DNA sequences and B) taxonomically-clustered pollen ASVs across model assessment metrics. Each axis on the radar plot is normalized to the minimum and maximum values across models, so that values at the center of the plot represent the minimum value across all models, and values at the edges represent the maximum. Moving clockwise from the top of the chart, metrics shown are model *R*^2^, average distance loss in kilometers, median absolute error (MAE), and root mean squared error (RMSE). Higher *R*^2^ values are preferable, while lower average distance loss, MAE, and RMSE are preferred when considering model performance. The latter three metrics were plotted as absolute values, so that the models covering the largest space in the radar chart correspond to the best-performing models across all metrics.

**Figure 2:**
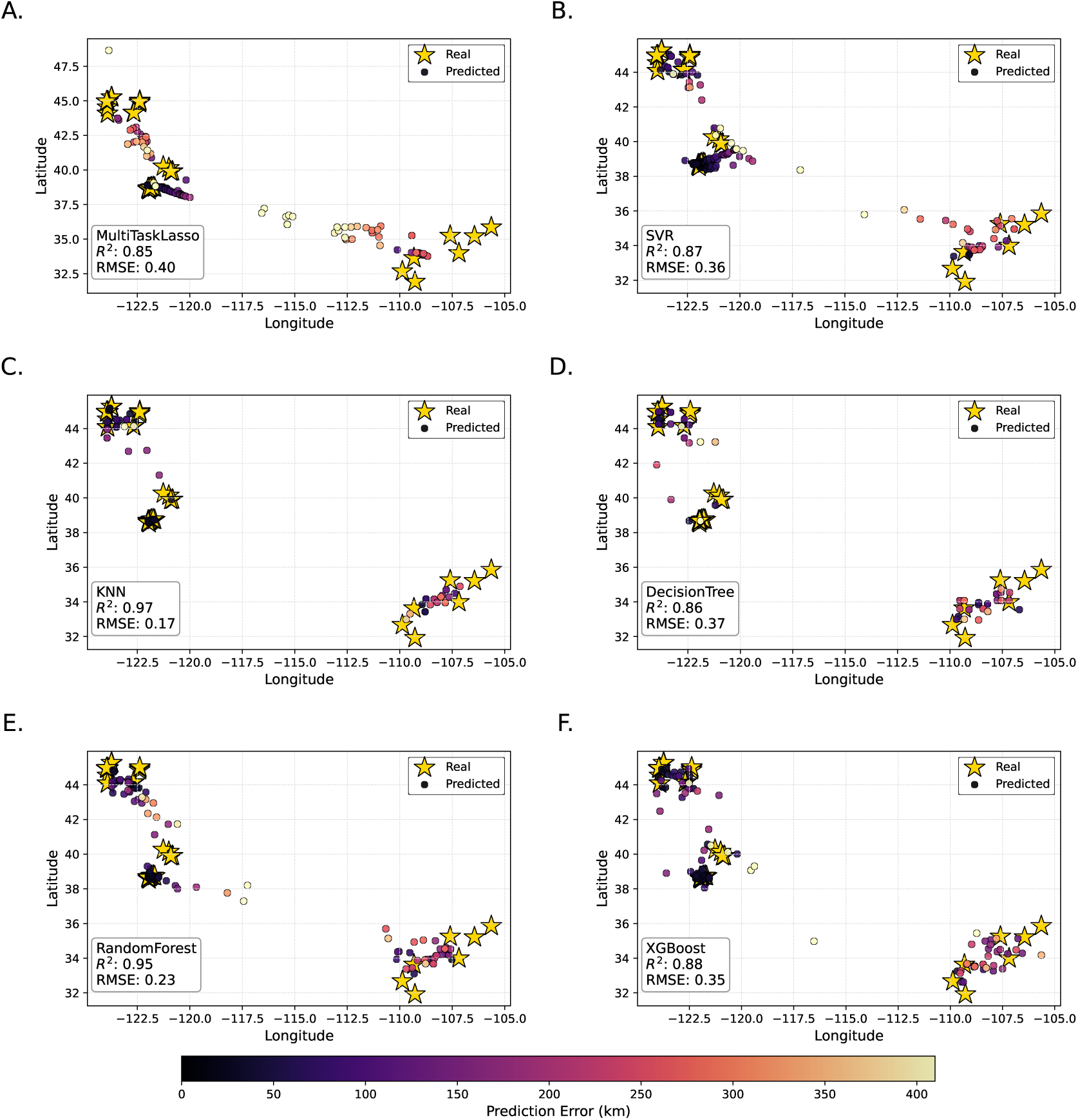
Location predictions from multioutput machine learning models trained on taxonomically clustered pollen ASVs. Yellow stars indicate true sample locations; predicted locations are shown as points colored by geodesic distance error (brighter colors indicate higher error). Inset boxes display model name, *R*^2^, and root mean squared error. Panels A–D show base model predictions: A) MultiTaskLasso, B) Support Vector Regressor, C) k-Nearest Neighbors, D) Decision Tree. Panels E–F show ensemble model predictions: E) XGBoost, F) Random Forest. Site clusters correspond to the three study areas moving from northwest to southeast: PNW Forests, CA Sunflowers, and Sky Islands.

**Figure 3:**
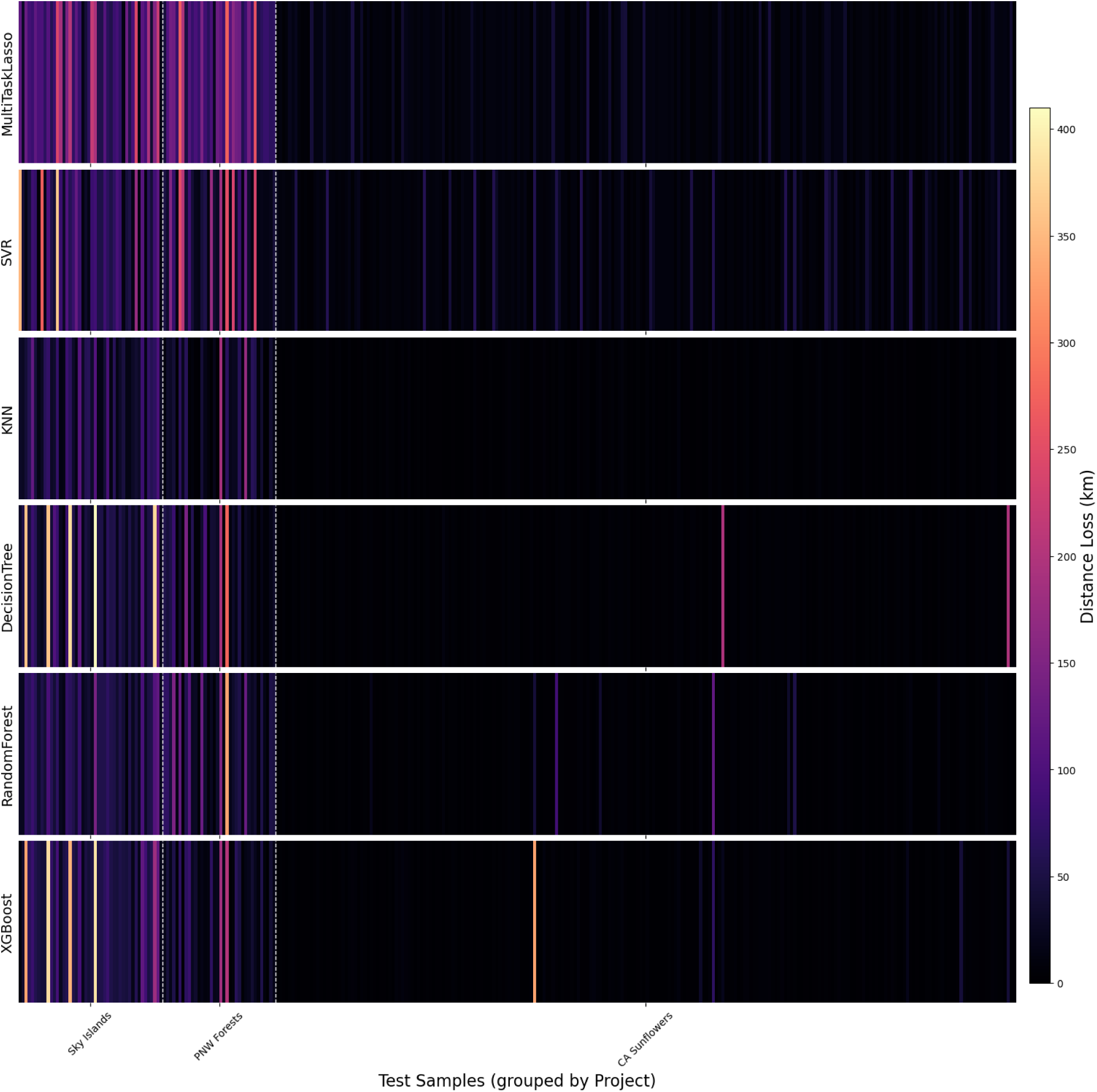
Distance error (km) for each test sample across models trained on taxonomically clustered pollen ASVs, calculated as the geodesic distance between true and predicted locations. Darker colors indicate lower error; brighter colors indicate higher error. Each vertical bar represents one sample, stacked to enable model comparison. Vertical dashed white lines separate projects, labeled on the x-axis. Sample IDs are omitted to avoid overcrowding.

**Figure 4:**
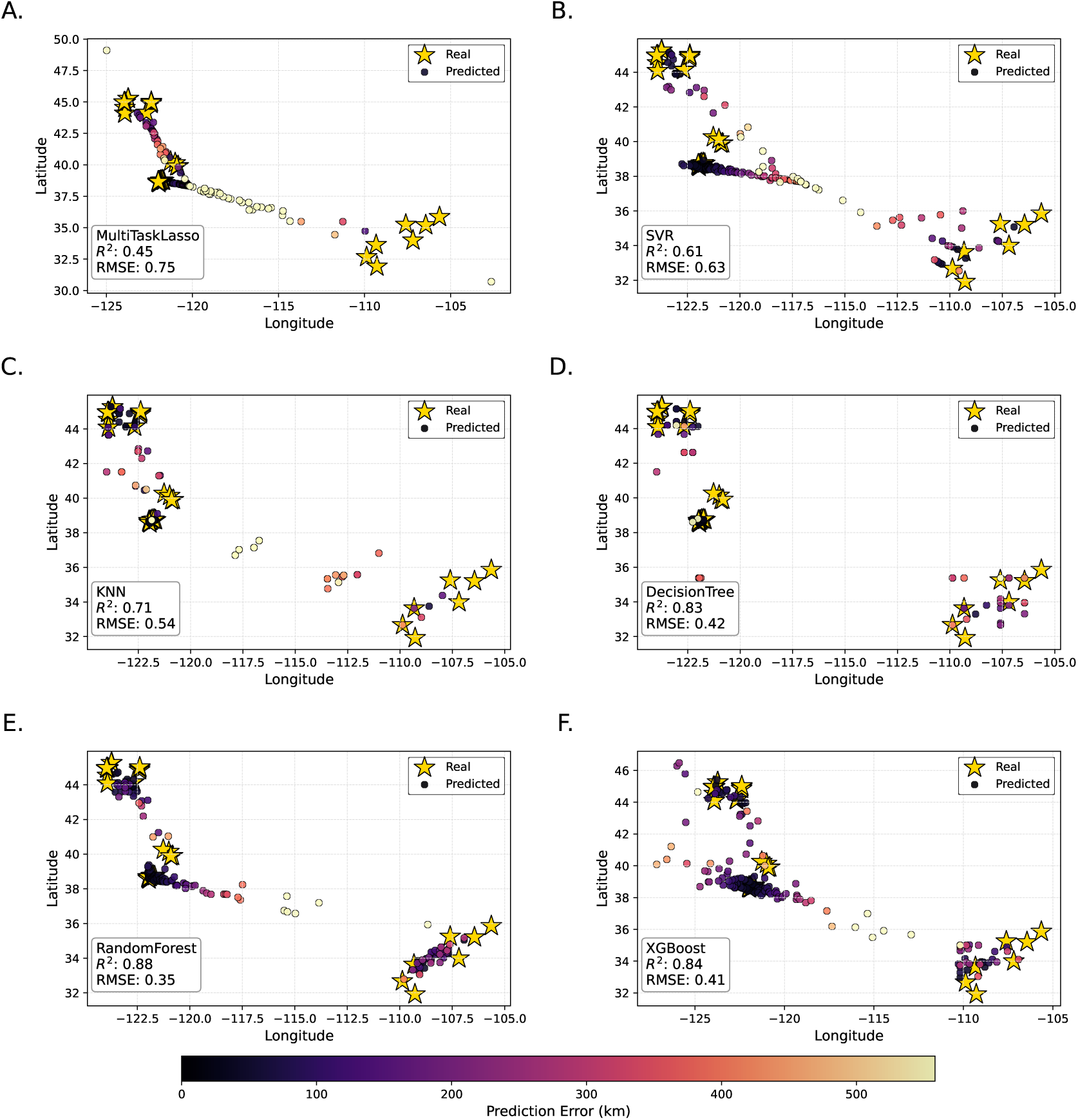
Location predictions from multioutput machine learning models trained on raw DNA sequences. Yellow stars mark true sample locations; predicted locations are shown as points colored by geodesic distance error (brighter colors indicate higher error). Inset boxes display model name, *R*^2^, and root mean squared error. Panels A–D show base models: A) MultiTaskLasso, B) Support Vector Regressor, C) k-Nearest Neighbors, D) Decision Tree. Panels E–F show ensemble models: E) XGBoost, F) Random Forest. Site clusters correspond to the three study areas moving from northwest to southeast: PNW Forests, CA Sunflowers, and Sky Islands.

**Figure 5:**
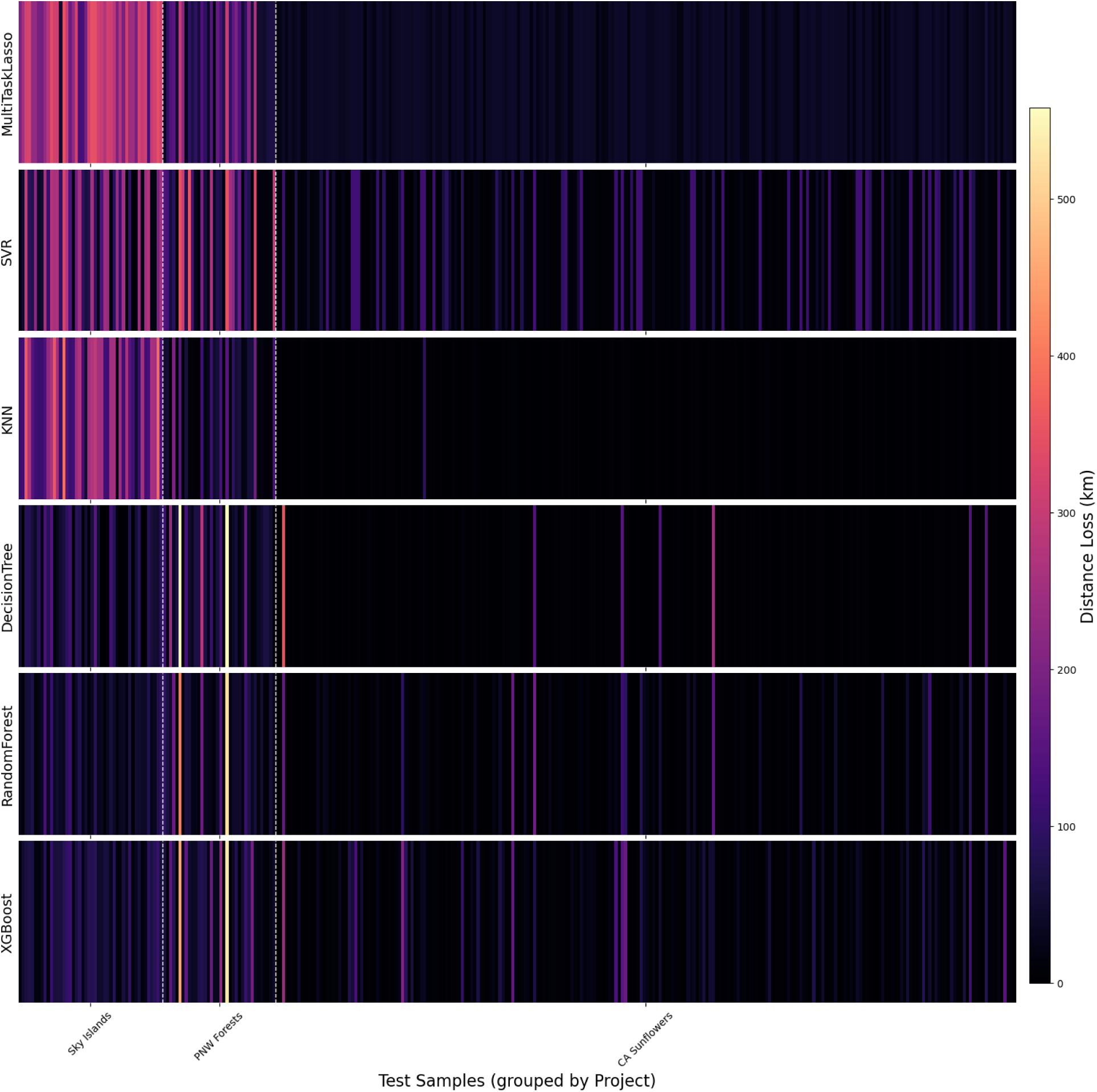
Distance error (km) for each test sample across models trained on raw pollen DNA sequences, calculated as the geodesic distance between true and predicted locations. Darker colors indicate lower error; brighter colors indicate higher error. Each vertical bar represents a single sample, aligned vertically to allow model comparison. Vertical dashed white lines separate projects, labeled on the x-axis. Sample IDs are omitted to avoid overcrowding.

**Table 1:** Performance metrics for best-tuned models across datasets trained on taxonomically-classified pollen sequence data, including *R*^2^, root mean squared error, median absolute error, average distance loss in kilometers with standard deviation.

| ALL | $R^2$ | RMSE | MAE | Avg. Distance Loss | Standard Deviation |
| --- | --- | --- | --- | --- | --- |
| DecisionTree | 0.8668 | 0.3736 | 0.0232 | 19.21 | 55.39 |
| KNN | 0.9761 | 0.1525 | 0.0156 | 10.16 | 22.89 |
| MultiTaskLasso | 0.8490 | 0.3973 | 0.0748 | 36.15 | 51.14 |
| RandomForest | 0.9507 | 0.2235 | 0.0202 | 15.65 | 32.29 |
| SVR | 0.8817 | 0.3503 | 0.0718 | 27.70 | 48.09 |
| XGBoost | 0.9211 | 0.2861 | 0.0204 | 17.24 | 41.91 |
| **California Sunflowers** |  |  |  |  |  |
| DecisionTree | -17.5831 | 0.0941 | 0.0227 | 5.18 | 17.89 |
| KNN | 0.1863 | 0.0222 | 0.0127 | 2.83 | 2.12 |
| MultiTaskLasso | -17.3350 | 0.0966 | 0.0635 | 12.55 | 8.86 |
| RandomForest | -9.2497 | 0.0698 | 0.0140 | 4.51 | 10.35 |
| SVR | -27.0492 | 0.1237 | 0.0597 | 13.80 | 13.70 |
| XGBoost | -2.8632 | 0.0465 | 0.0156 | 4.08 | 7.25 |
| **Pacific Northwest Forests** |  |  |  |  |  |
| DecisionTree | 0.1287 | 0.3835 | 0.0924 | 41.08 | 56.44 |
| KNN | 0.4603 | 0.2963 | 0.0913 | 33.28 | 41.48 |
| MultiTaskLasso | -1.3819 | 0.6590 | 0.4694 | 109.63 | 59.24 |
| RandomForest | -0.7685 | 0.4854 | 0.1829 | 54.80 | 61.55 |
| SVR | -1.5536 | 0.6194 | 0.2227 | 74.81 | 78.49 |
| XGBoost | 0.0846 | 0.3961 | 0.1451 | 49.07 | 53.44 |
| **Sky Islands** |  |  |  |  |  |
| DecisionTree | -6.8594 | 0.8465 | 0.1829 | 74.05 | 112.62 |
| KNN | 0.2623 | 0.2866 | 0.0851 | 29.68 | 34.29 |
| MultiTaskLasso | -5.0615 | 0.7314 | 0.5393 | 99.69 | 61.72 |
| RandomForest | -0.1153 | 0.3449 | 0.2112 | 42.14 | 34.01 |
| SVR | -3.1133 | 0.6410 | 0.1530 | 62.10 | 79.32 |
| XGBoost | -2.8124 | 0.6285 | 0.2246 | 59.89 | 78.33 |

**Table 2:**
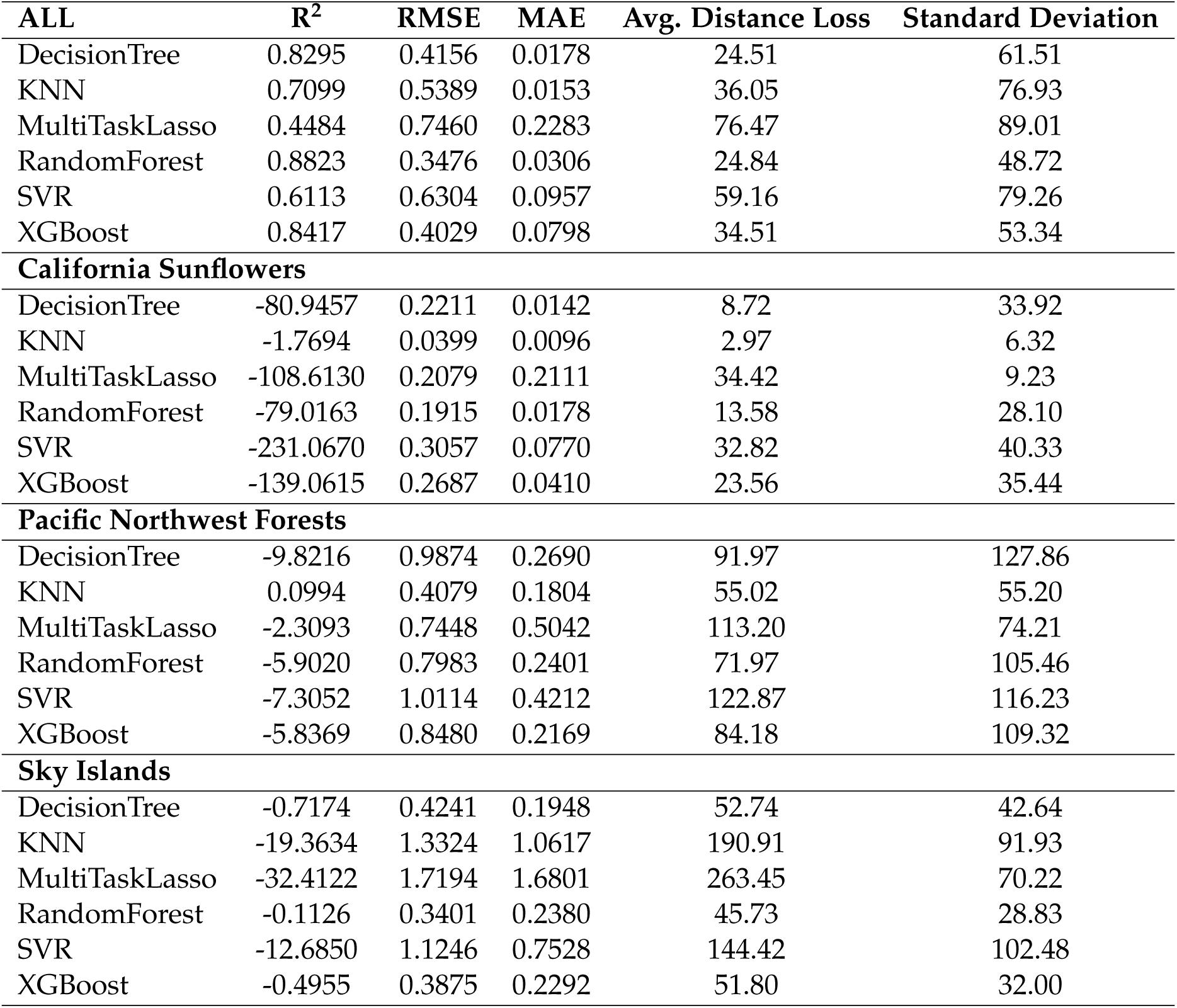
Performance metrics for best-tuned models across datasets trained on raw pollen sequence data, including R^2^, root mean squared error, median absolute error, average distance loss in kilometers with standard deviation.

| **ALL** | **$R^2$** | **RMSE** | **MAE** | **Avg. Distance Loss** | **Standard Deviation** |
| --- | --- | --- | --- | --- | --- |
| DecisionTree | 0.8295 | 0.4156 | 0.0178 | 24.51 | 61.51 |
| KNN | 0.7099 | 0.5389 | 0.0153 | 36.05 | 76.93 |
| MultiTaskLasso | 0.4484 | 0.7460 | 0.2283 | 76.47 | 89.01 |
| RandomForest | 0.8823 | 0.3476 | 0.0306 | 24.84 | 48.72 |
| SVR | 0.6113 | 0.6304 | 0.0957 | 59.16 | 79.26 |
| XGBoost | 0.8417 | 0.4029 | 0.0798 | 34.51 | 53.34 |
| **California Sunflowers** |  |  |  |  |  |
| DecisionTree | -80.9457 | 0.2211 | 0.0142 | 8.72 | 33.92 |
| KNN | -1.7694 | 0.0399 | 0.0096 | 2.97 | 6.32 |
| MultiTaskLasso | -108.6130 | 0.2079 | 0.2111 | 34.42 | 9.23 |
| RandomForest | -79.0163 | 0.1915 | 0.0178 | 13.58 | 28.10 |
| SVR | -231.0670 | 0.3057 | 0.0770 | 32.82 | 40.33 |
| XGBoost | -139.0615 | 0.2687 | 0.0410 | 23.56 | 35.44 |
| **Pacific Northwest Forests** |  |  |  |  |  |
| DecisionTree | -9.8216 | 0.9874 | 0.2690 | 91.97 | 127.86 |
| KNN | 0.0994 | 0.4079 | 0.1804 | 55.02 | 55.20 |
| MultiTaskLasso | -2.3093 | 0.7448 | 0.5042 | 113.20 | 74.21 |
| RandomForest | -5.9020 | 0.7983 | 0.2401 | 71.97 | 105.46 |
| SVR | -7.3052 | 1.0114 | 0.4212 | 122.87 | 116.23 |
| XGBoost | -5.8369 | 0.8480 | 0.2169 | 84.18 | 109.32 |
| **Sky Islands** |  |  |  |  |  |
| DecisionTree | -0.7174 | 0.4241 | 0.1948 | 52.74 | 42.64 |
| KNN | -19.3634 | 1.3324 | 1.0617 | 190.91 | 91.93 |
| MultiTaskLasso | -32.4122 | 1.7194 | 1.6801 | 263.45 | 70.22 |
| RandomForest | -0.1126 | 0.3401 | 0.2380 | 45.73 | 28.83 |
| SVR | -12.6850 | 1.1246 | 0.7528 | 144.42 | 102.48 |
| XGBoost | -0.4955 | 0.3875 | 0.2292 | 51.80 | 32.00 |

The Random Forest model yielded the best fit for the raw pollen ASVs and the second-best fit for the taxonomically-clustered pollen ASVs (Table 1, Table 2, Fig. 1). Since the random forest performed well across both training datasets, we investigated feature importance to understand which raw and taxonomically clustered pollen ASVs drove predictions. Because we predicted latitude and longitude separately as a multioutput regression, there are separate feature importances for each (Figs. S 5 and 6). After determining the top 25 feature importances for both the taxonomic and raw sequence datasets, we used NCBI BLAST on the high-importance raw sequences. We then assigned the genus and species from the top hit to compare which plant species drove predictions for both training datasets (Fig S 5).

We found that plants from the genera *Rubus*, *Helianthus*, *Digitalis*, *Gaultheria*, and *Phacelia* were particularly important for latitude predictions across both the raw and the taxonomic-clustered training sets. For longitude predictions, multiple features that matched to the families Asteraceae, Brassicaceae, Rosaceae, and Fabaceae were highly important. The taxonomic-clustered training set was often matched to a lower resolution due to our cross-referencing of NCBI and RDP; however, it returned several highly reliable species-level features that were highly important for predictions. These included *Phacelia heterophylla*, *Helianthus annuus*, *Taraxacum officinale*, *Vicia americana*, *Campanula rotundifolia*, *Hypericum elodeoides*, *Cichorium intybus*, *Convolvulus arvensis*, *Daucus carota*, *Erigeron canadensis*, *Brassica rapa*, and *Capsella bursa-pastoris*.

## 4 Discussion

In this study, we demonstrated the potential of machine learning models to predict the geographic origin of mixed-species pollen samples collected by bees, providing a practical framework for integrating pollen DNA sequencing with machine learning for geolocation tasks. Across all models tested, ensemble approaches consistently achieved the highest geolocation accuracy while minimizing error, as observed in multiple contexts (e.g. Zhang and Ma, 2012). However, since each model type carries distinct trade-offs, comparing predictions across multiple algorithms remains an important strategy to maximize overall predictive performance.

We also evaluated how the format of the pollen data — whether taxonomically classified or raw DNA sequences — affected model performance. All models performed better when trained on taxonomically classified ASVs, but ensemble models and k-Nearest Neighbors algorithms maintained strong performance across both data types. The taxonomically classified dataset, consisting of 185 unique pollen features, reflects a more curated approach in which sequence variants were clustered into species. In contrast, the raw DNA sequence dataset included 954 unique features, with each sequence variant treated as a distinct taxon. While the taxonomic classification process helps simplify the dataset, it introduces potential bias through manual assignments. Using raw DNA sequences offers a less biased alternative and better reflects the data typically available in public DNA repositories, making it a more realistic option for geolocation applications. In this workflow, once a predictive model is trained on raw sequences and the most informative features are identified, these features can be matched to BLAST databases to obtain taxonomic identities, if needed. This approach not only aids in sample geolocation but also highlights which plant species may hold the most predictive value. Overall, our findings position machine learning on pollen DNA barcoding data as a powerful new tool for palynology.

Across multiple projects, we found that poor predictive performance was often associated with pollen samples that contained limited geographic information. For example, in the Sky Islands project, several poorly predicted samples were dominated by a single pollen type identified only at the family level, such as Asteraceae or Rosaceae. These families are both taxonomically diverse and geographically widespread, spanning the entire spatial extent of the study region. Similarly, in the California Sunflower project, the least accurately predicted samples were composed primarily of widespread, weedy species (e.g., *Taraxacum officinale*, *Achillea millefolium*, *Erigeron canadensis*, *Cicuta maculata*, and *Cichorium intybus*). Many of these taxa are not morphologically distinct, highlighting the value of using DNA sequence data: *Taraxacum* and *Cichorium* are both in the subfamily Cichorioideae, while *Erigeron* and *Achillea* are both in the subfamily Asteroideae, each of which are typically the finest taxonomic resolution that can be visually distinguished (Erdtman, 2023). The presence of such broadly distributed taxa — whether due to low taxonomic resolution or ecological ubiquity — provides little discriminatory power for geolocation tasks, as these species can be found across much of the landscape. These findings suggest that future models may benefit from training datasets that exclude widespread or ecologically ubiquitous species, and, if possible, focus instead on more range-restricted and geographically informative taxa. Previous work has proposed classifications of pollen types based on their forensic value, emphasizing rare or endemic species as more informative for geolocation; however, such plants are also less likely to be encountered in environmental samples, creating a trade-off between informativeness and detectability (Helderop et al., 2021). Integrating such classifications into the geolocation workflow could help differentiate between samples that lack predictive accuracy due to ecological limitations and those where the model itself has failed.

In addition to samples with low-information pollen profiles, model performance was also limited at sites with sparse training data. For the most poorly predicted samples from the Pacific Northwest Forests project, the low accuracy appeared to stem from insufficient representation in the training dataset. For example, the two sites with the lowest predictive accuracy included fewer than 5 samples across both the training and test datasets. Furthermore, the limited training samples from these sites showed considerable variability in pollen assemblage, providing the models with inconsistent and weakly location-specific examples from which to learn. Class imbalance has been a ubiquitous challenge for machine learning practitioners in both classification and regression tasks (Branco et al., 2016), and these findings suggest that adequate sample representation is critical for reliable geographic prediction. Future studies could improve model performance by establishing a minimum sample threshold for including a site or project in the training dataset, balancing the trade-off between geographic coverage and predictive accuracy.

When evaluated on each project individually, several models produced negative R^2^ values, indicating performance worse than a baseline model predicting the project mean. This does not imply model failure per se; rather, it reflects substantial distributional differences among projects. Because models were trained on the combined dataset, project-specific patterns in ASV composition and the taxonomic–geographic relationships were not fully captured, reducing transferability across regions. Negative R^2^ values therefore highlight strong biogeographic turnover and region-specific community structure that limit cross-project generalization. Still, for the goal of large-scale geolocation tasks, samples from a wide spatial distribution are required and thus pulling from multiple project sources will continue to be necessary.

Building on the observation that both taxonomic resolution and the representation of training data affect model performance, we compared feature importance across taxonomically clustered and raw ASV training datasets to assess whether key predictive signals were preserved. In many cases, the underlying drivers of model performance appeared consistent across data types. For instance, in the clustered dataset, taxa matching the genus *Rubus* (e.g., blackberries) emerged as highly influential for geolocation. This pattern was mirrored in the raw ASV dataset, where multiple distinct *Rubus* sequence variants also ranked highly in importance, suggesting that species-level diversity within *Rubus* contributes valuable geographic signal across the study regions. A similar pattern was observed for the family Ericaceae, which ranked highly in the clustered dataset, while raw ASVs identified as multiple *Rhododendron* species were prominent in the unclustered data. These consistencies suggest that, despite modest improvements in predictive accuracy when using the taxonomically clustered dataset, the core spatial signals are retained in the raw sequence data. Consequently, the additional effort required for taxonomic clustering — both computational and manual curation — may not be necessary for all applications. Ultimately, the decision to cluster sequences before modeling can be guided by each practitioner’s specific goals, resources, and downstream needs.

An additional consideration in evaluating model performance is the spatial resolution achievable with these models. While our approach proved effective for distinguishing among broad regions and states, its accuracy diminished at finer spatial resolutions. The appropriate level of resolution will depend on the specific goals of an investigation. In some historical forensic cases, determining the state or country of origin was sufficient for drawing meaningful conclusions (Bryant and Jones, 2006). However, there are likely scenarios — such as identifying the provenance of illicit goods or tracking pollinator movements — where finer-scale resolution would be beneficial or even necessary. Expanding the training dataset, both in size and geographic breadth, could help achieve this. It is common in supervised machine learning to provide training datasets of thousands or more samples; thus, our dataset represents the lower end of sample size for model training. Moreover, integrating additional data sources — such as species distribution models, climate variables, geological information, or other ecological layers — could further refine spatial predictions. Furthermore, beyond incorporating more data, combining multiple techniques to optimize accuracy and minimize computational strain has been previously proposed and has potential to facilitate even better predictions. For example, utilizing species distribution models to enable network-based identification of impossible sites of origin can serve as a ‘first-pass filter’ to create a smaller training dataset with more explanatory power (Helderop et al., 2021). This refined training data can then be passed to more computationally intensive geoforensic models, such as the Geoforensic Interdiction (GOFIND) model, which can successfully identify multiple potential origin sites from USDA CropScape data while avoiding predictions at impossible sites (Tong et al., 2021). Our modeling framework utilizes known pollen assemblage data from bee-collected pollen, which could provide an additional validation step in such a pipeline: impossible locations could be filtered using network analysis, multiple possible locations of origin could be identified using the GOFIND model, then bee-collected pollen data could be used to further refine predictions using actual ground-truthed pollen data rather than just relying on possible species distributions. As the field of palynology grows and more tools and analytical techniques are developed, it remains important to consider how they can work together to achieve more reliable inference. Given the flexibility of machine learning algorithms to incorporate diverse types of input data, there is considerable potential to build more powerful and nuanced geolocation tools. Nonetheless, our results show that accurate predictions within tens of kilometers are already possible using relatively modest amounts of pollen DNA sequence data alone, establishing a strong foundation for future methodological advances.

Although this study focuses on metabarcoded pollen DNA from bees to demonstrate the potential of machine learning algorithms for geographic prediction, the underlying framework is broadly applicable to a range of sample types comprising DNA from multiple species. It was previously established that sample geolocation can be achieved using machine learning algorithms trained on individual whole-genome sequences (Battey et al., 2020), but here we show that short metabarcoded sequences in multispecies samples can also produce reliable location predictions. Compared to whole-genome sequencing, metabarcoding is cheaper and faster, with more publicly available reference libraries, and thus represents an attractive alternative that still produces accurate predictions. Despite the limitations of current pollen reference libraries, sequences derived from airborne monitoring programs, sediment cores used in paleopalynological research, or even historical samples from museum collections could all contribute to the development of expanded reference libraries for spatial inference. While we centered our analyses on the rbcL gene — commonly used in pollinator diet studies — other conserved plant barcoding markers, such as matK, could be readily incorporated into the same modeling pipeline. Expanding the number and type of genetic markers would enhance taxonomic resolution and increase the breadth of reference datasets. More broadly, this study presents a generalizable workflow for cleaning sequence data and training machine learning models for geolocation tasks. Future practitioners are encouraged to adapt this framework to meet the specific requirements of their research questions, incorporating additional data sources as needed to improve resolution, accuracy, and ecological relevance.

Our results underscore the value of repurposing existing pollen metabarcoding data for geolocation applications. A central challenge in advancing palynology has been the scarcity of reference pollen libraries, which are essential for accurate spatial prediction (Bryant and Jones, 2006). This shortage has likely created a self-reinforcing barrier, where the high cost and expertise required to collect and curate reference datasets — particularly those based on morphological identification — have limited progress to a few specialized institutions. By contrast, our workflow demonstrates that accurate geolocation is achievable using raw pollen DNA sequences, bypassing the need for expert morphological identification. Taxonomic assignments can still be performed bioinformatically when needed, offering flexibility without compromising accessibility. While machine learning approaches have not yet been widely adopted within palynology, this study illustrates their substantial potential: models trained on existing metabarcoding datasets can effectively recover spatial information, reducing reliance on traditional, labor-intensive methods. By lowering the technical and resource barriers, our approach opens new pathways for expanding the reach and utility of palynology, enabling broader adoption across diverse research and investigative contexts.

## 4.1 Acknowledgements

We extend our heartfelt gratitude to the more than 50 field crew members who dedicated their time and effort to sample collection across all three projects—this research would not have been possible without their commitment and hard work. We thank Jocelyn Zorn, Hamutahl Cohen, and Gordon Smith for their valuable contributions to the molecular work on the Sky Islands and California Sunflower projects, and H. Cohen, G. Smith, and Laura Jones for essential bioinformatics support. Field crew leadership was expertly provided by H. Cohen, G. Smith, Kaysee Arrowsmith, Jess Mullins, Jesse Fan Brown, Claire Massaro, Felix Bruner, Kylie Weeks, and Rose McDonald. The Sky Islands project received funding through NSF PCE Award 2009075 to LCP and Shalene Jha. The Pacific Northwest Forests project was supported by USDA AFRI Award 2023-67013-39910 to LCP, Katie Moriarty, Jim Rivers, and Lauren Grand. Additional funding was provided by the National Council for Air and Stream Improvement, Inc. (NCASI) and by the NCASI Foundation, supporting grants to K. Moriarty from the US Fish and Wildlife Service’s Wildlife Conservation Initiative, facilitated by Vicki Finn and Matt Dekar, and USDA Forest Service Agreements 20-PA-11061200-013 and 22-CS-11221632-029 facilitated by Deanna Williams. The California Sunflowers project was funded through the Foundation for Food and Agriculture grant CA18-SS-0000000009 to LCP, Quinn McFrederick, and Hollis Woodard. We particularly acknowledge NCASI’s pivotal role in making the Pacific Northwest Forests project possible through their collaboration and resources, the landowner’s participation, and the coordination and leadership from K. Moriarty for study design and data collection efforts. RAH received support through NSF GRFP Award 2137533. ADK was funded in part by NIH awards R01HG010774 and R35GM148253. Finally, we are grateful to the sunflower growers in California’s Central Valley, private forest owners in the Pacific Northwest, and the Siuslaw, Apache-Sitgreaves, Cibola, Coronado, Lincoln, and Santa Fe National Forests for providing access to research sites.

## 4.2 Author Contributions

RAH: Conceptualization, Methodology, Formal analysis, Investigation, Data Curation, Visualization, Writing - Original Draft, Writing - Review and Editing. ADK: Conceptualization, Methodology, Writing - Review and Editing. LCP: Conceptualization, Methodology, Data Curation, Supervision, Project administration, Writing - Review and Editing.

## 4.3 Data Accessibility

Raw data, novel code, and data products necessary for reproducing all analyses are deposited on Github (https://github.com/hayesrebecca/pollenGeolocation). The code in these external repositories is complete to allow replication of the tables, graphs, and statistical analyses that are reported in the publication.

## Supplementary Information

**Figure 1:**
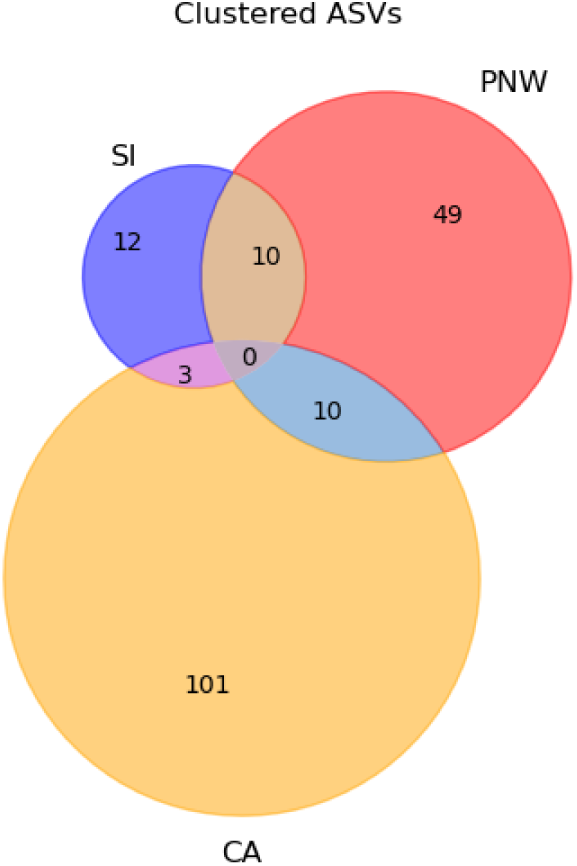
The overlap between projects of taxonomically clustered ASVs. Projects are encoded as follows: California Sunflowers Project (CA), Pacific Northwest Forests Project (PNW), and Sky Islands Project (SI).

**Figure 2:**
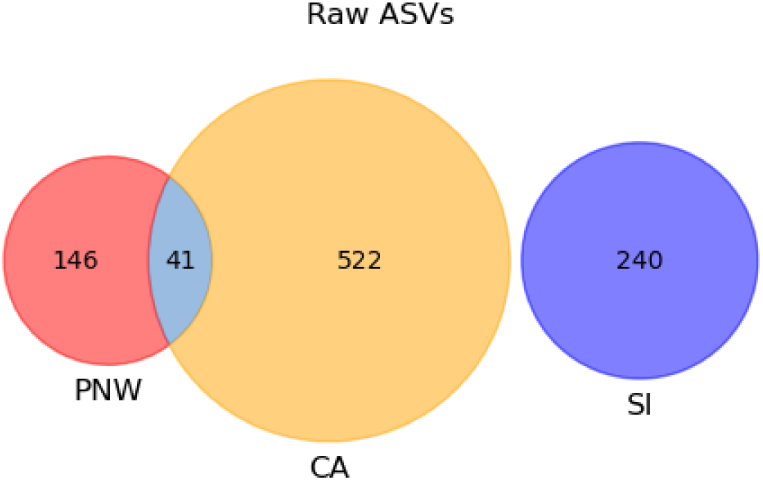
The overlap between projects of raw ASVs. Projects are encoded as follows: California Sunflowers Project (CA), Pacific Northwest Forests Project (PNW), and Sky Islands Project (SI).

**Table 1:**
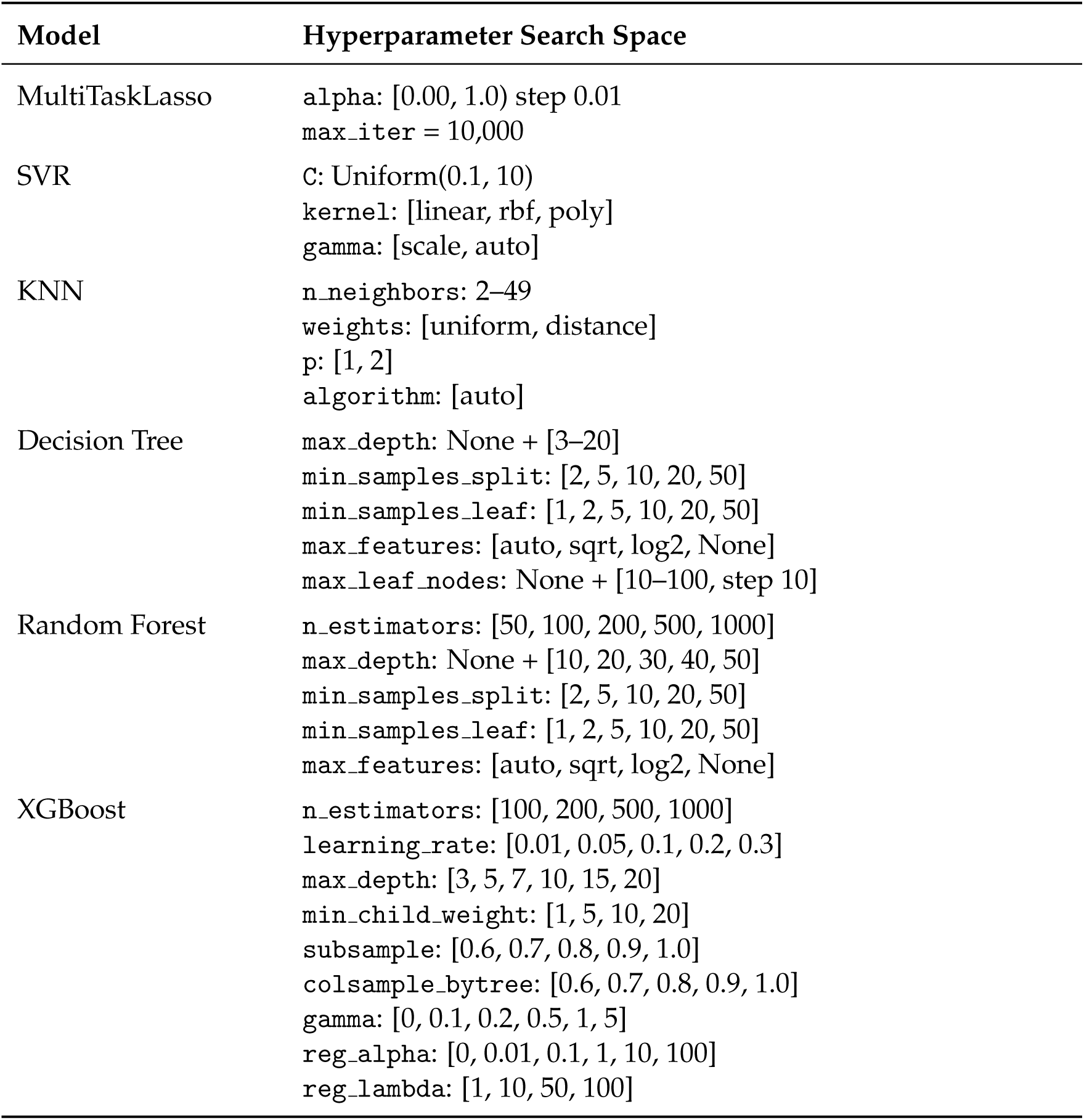
Hyperparameter search space for each model.

**Table 2:**
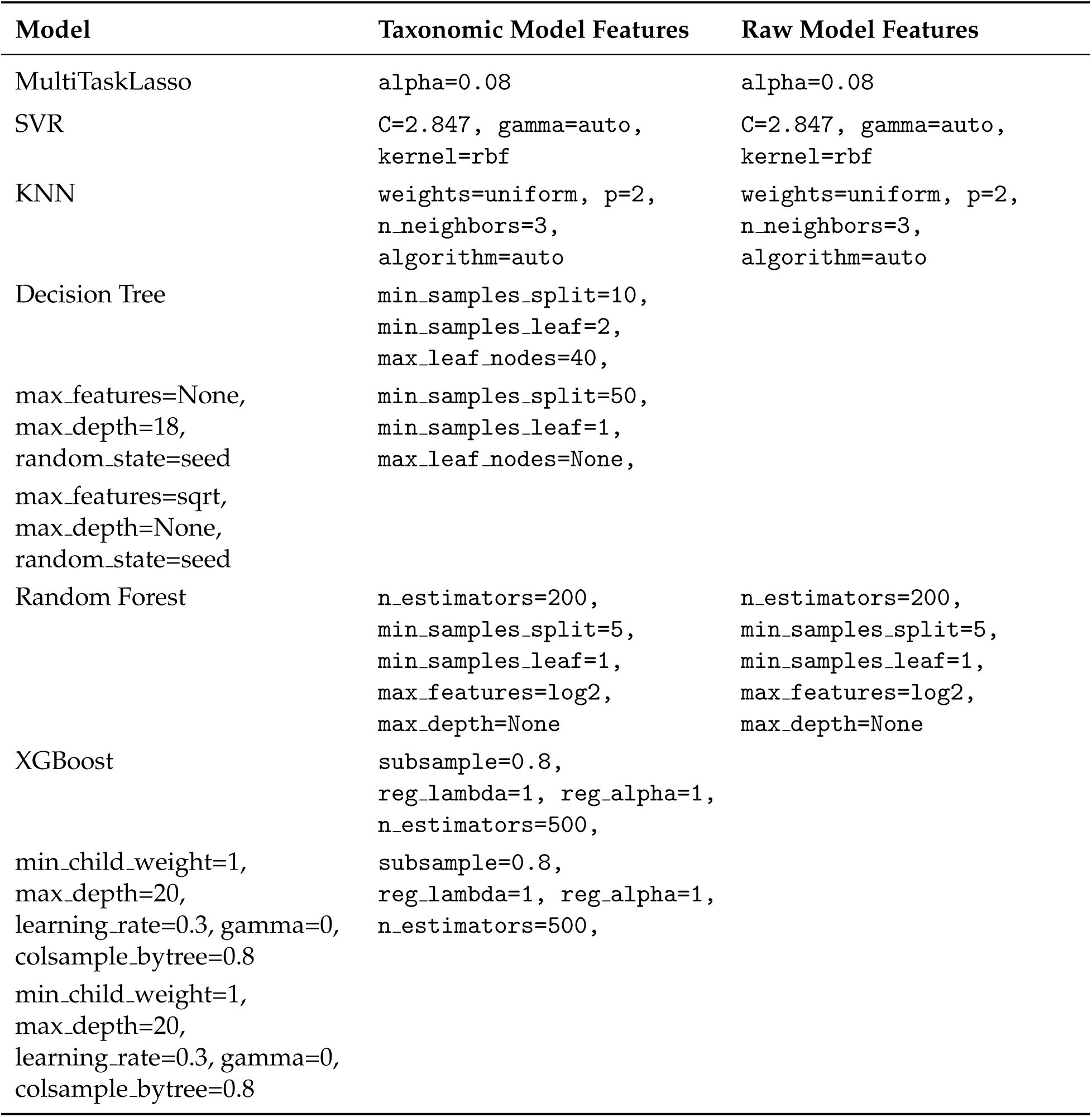
Best-tuned hyperparameters for taxonomic and raw models.

| Model | Taxonomic Model Features | Raw Model Features |
| --- | --- | --- |
| MultiTaskLasso | alpha=0.08 | alpha=0.08 |
| SVR | C=2.847, gamma=auto,<br>kernel=rbf | C=2.847, gamma=auto,<br>kernel=rbf |
| KNN | weights=uniform, p=2,<br>n_neighbors=3,<br>algorithm=auto | weights=uniform, p=2,<br>n_neighbors=3,<br>algorithm=auto |
| Decision Tree | min_samples_split=10,<br>min_samples_leaf=2,<br>max_leaf_nodes=40, |  |
| max_features=None,<br>max_depth=18,<br>random_state=seed | min_samples_split=50,<br>min_samples_leaf=1,<br>max_leaf_nodes=None, |  |
| max_features=sqrt,<br>max_depth=None,<br>random_state=seed |  |  |
| Random Forest | n_estimators=200,<br>min_samples_split=5,<br>min_samples_leaf=1,<br>max_features=log2,<br>max_depth=None | n_estimators=200,<br>min_samples_split=5,<br>min_samples_leaf=1,<br>max_features=log2,<br>max_depth=None |
| XGBoost | subsample=0.8,<br>reg_lambda=1, reg_alpha=1,<br>n_estimators=500, |  |
| min_child_weight=1,<br>max_depth=20,<br>learning_rate=0.3, gamma=0,<br>colsample_bytree=0.8 | subsample=0.8,<br>reg_lambda=1, reg_alpha=1,<br>n_estimators=500, |  |
| min_child_weight=1,<br>max_depth=20,<br>learning_rate=0.3, gamma=0,<br>colsample_bytree=0.8 |  |  |

**Figure 3:**
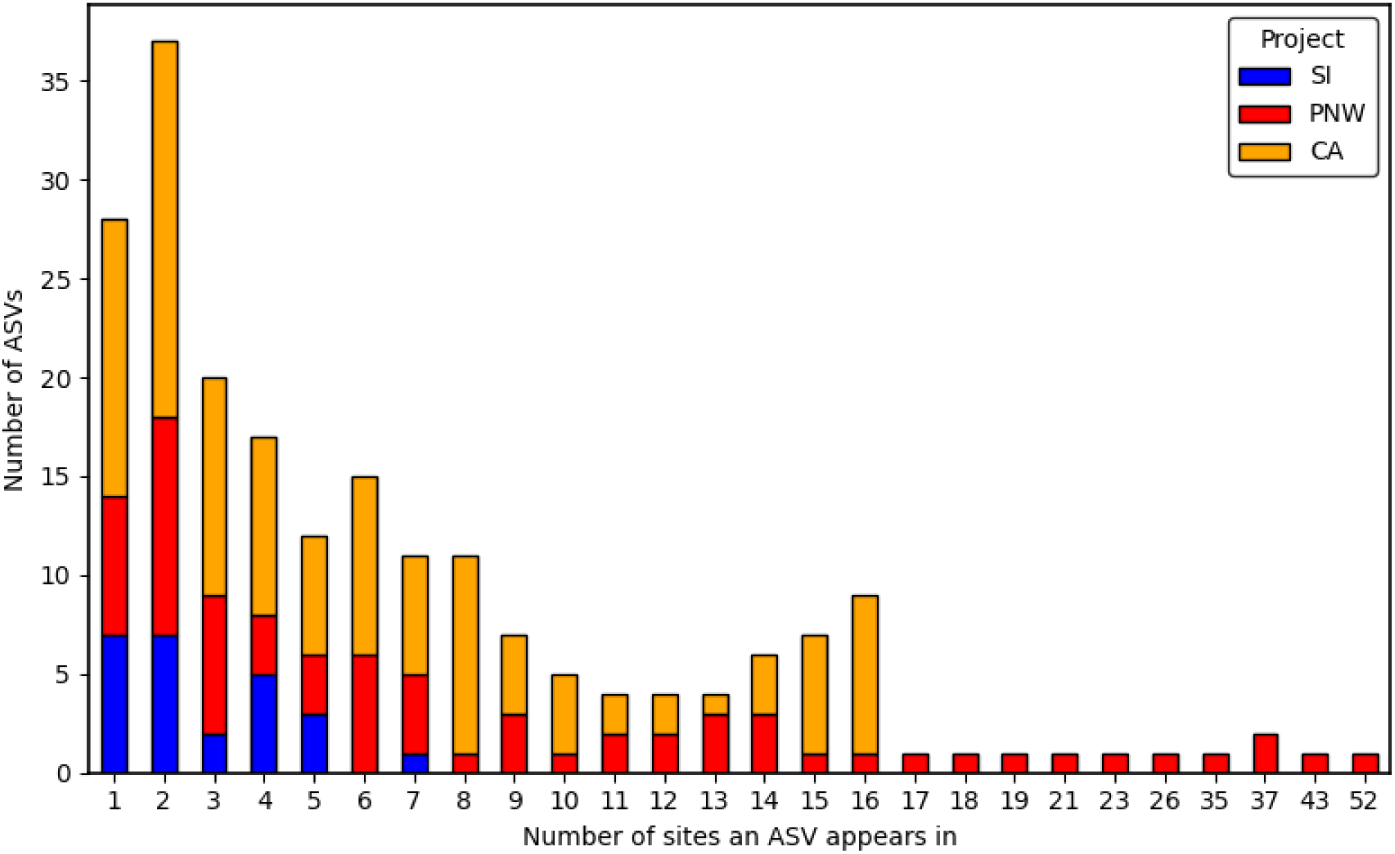
Distribution of taxonomically clustered ASV occurrence across sites by project. An ASV was considered present at a site if it was detected in at least one individual at a given site for that project. Projects are encoded as follows: California Sunflowers Project (CA), Pacific Northwest Forests Project (PNW), and Sky Islands Project (SI).

**Figure 4:**
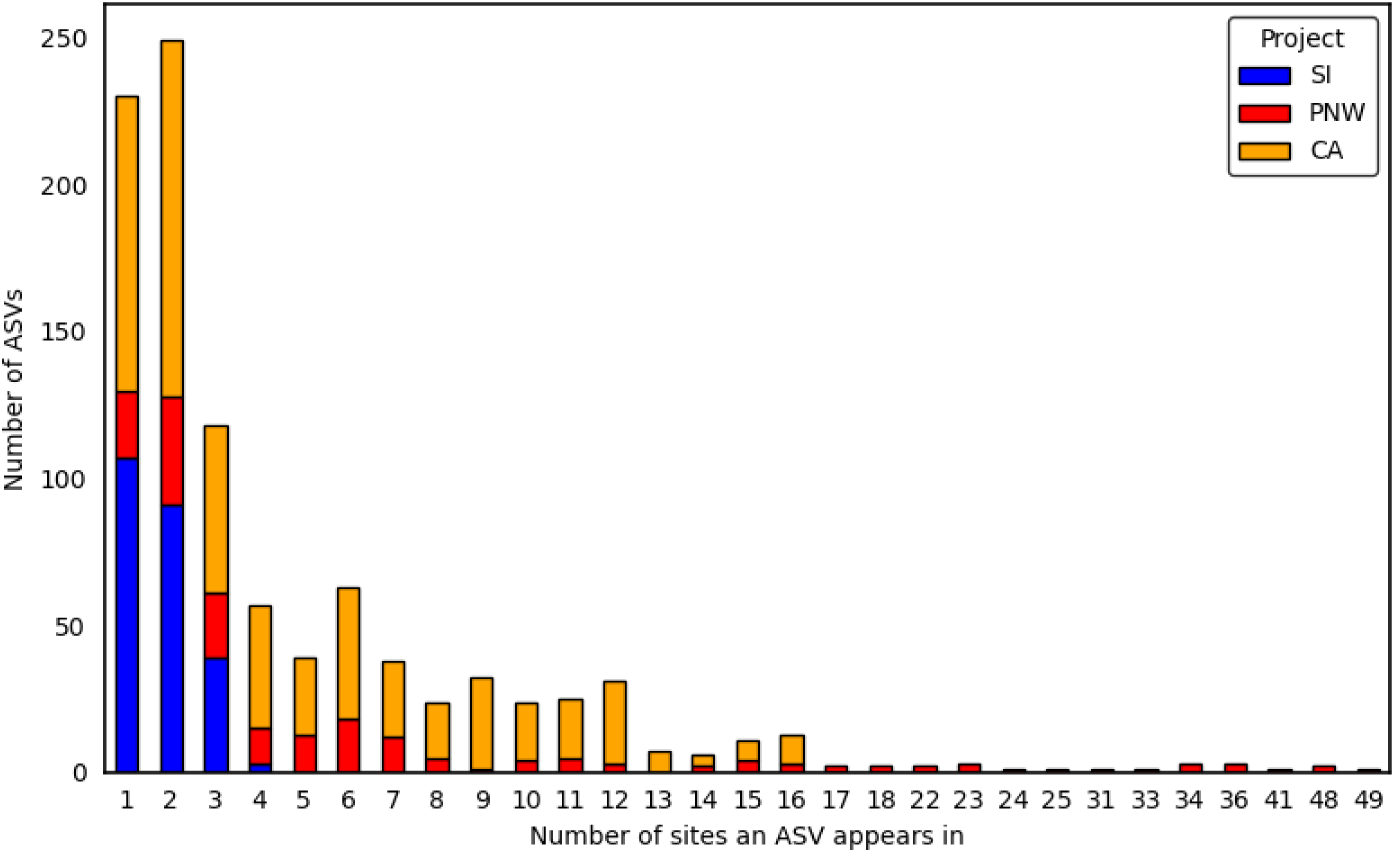
Distribution of raw ASV occurrence across sites by project. An ASV was considered present at a site if it was detected in at least one individual at a given site for that project. Projects are encoded as follows: California Sunflowers Project (CA), Pacific Northwest Forests Project (PNW), and Sky Islands Project (SI).

**Figure 5:**
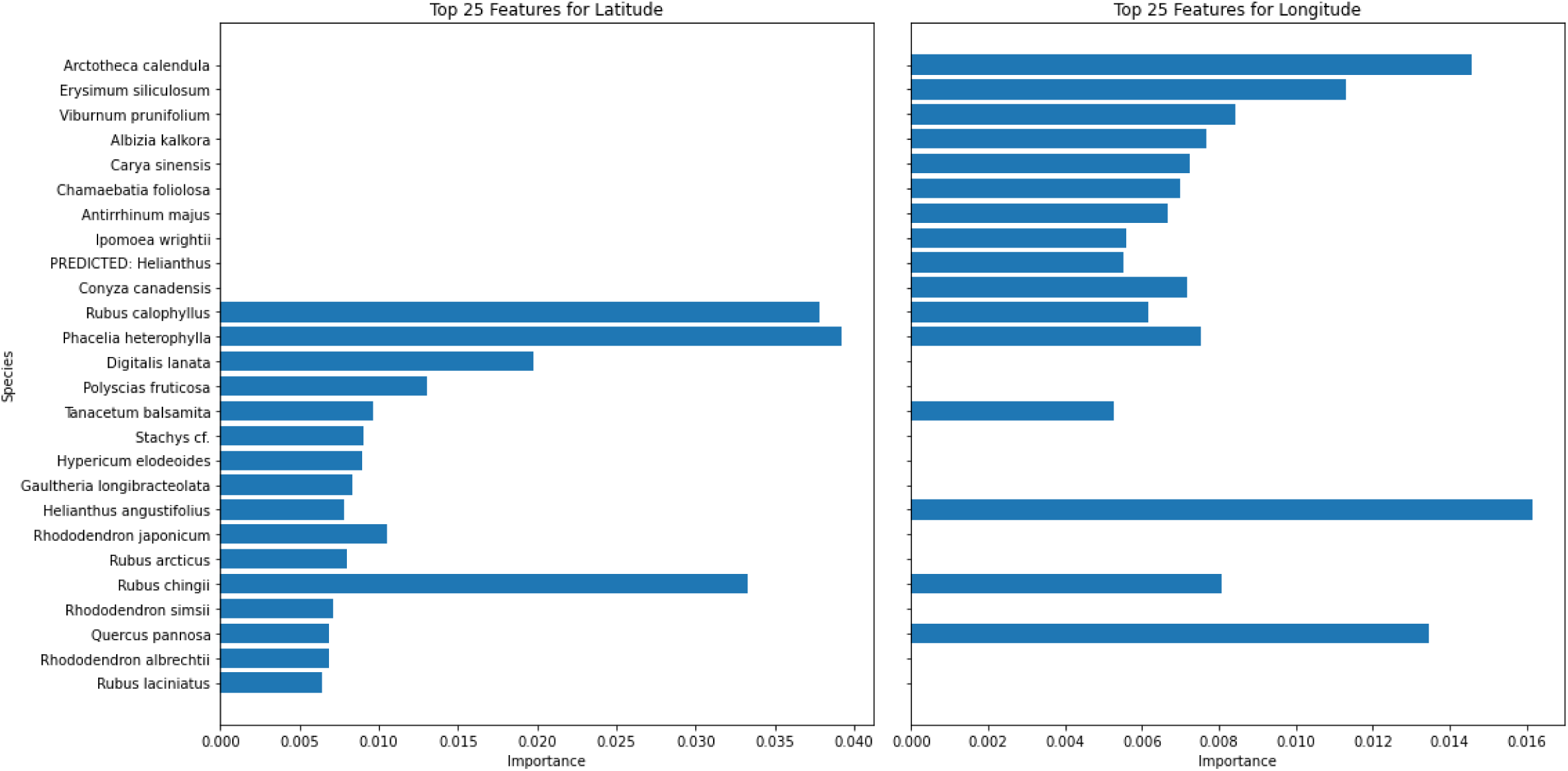
Feature importances for top 25 most informative ASVs for the models trained on raw sequence data. Feature importances were extracted and then classified taxonomically using NCBI BLAST.

**Figure 6:**
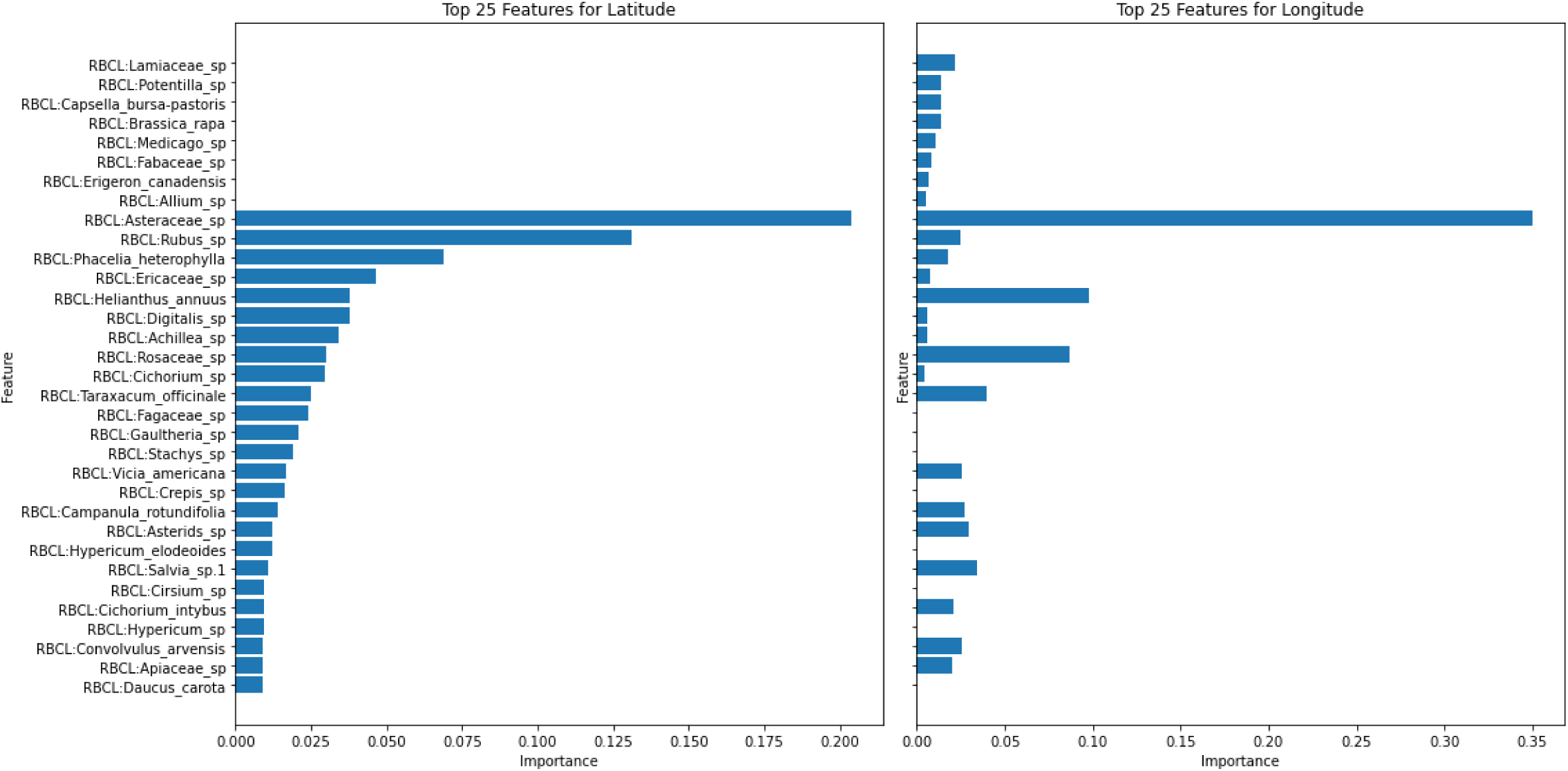
Feature importances for top 25 most informative ASVs for the models trained on taxonomically-assigned data.

## Notes

https://github.com/hayesrebecca/pollenGeolocation

### Competing Interest Statement

The authors have declared no competing interest.

## References

1, C. P. W. G., P. M. Hollingsworth, L. L. Forrest, J. L. Spouge, M. Hajibabaei, S. Ratnasingham, M. van der Bank, M. W. Chase, R. S. Cowan, D. L. Erickson, et al. 2009. A DNA barcode for land plants. Proceedings of the National Academy of Sciences 106:12794–12797.

Awad, M., R. Khanna, M. Awad, and R. Khanna. 2015. Support vector regression. Efficient learning machines: Theories, concepts, and applications for engineers and system designers pages 67–80.

Battey, C. J., P. L. Ralph, and A. D. Kern. 2020. Predicting geographic location from genetic variation with deep neural networks. Elife 9:e54507.

Bell, K. L., K. S. Burgess, J. C. Botsch, E. K. Dobbs, T. D. Read, and B. J. Brosi. 2019. Quantitative and qualitative assessment of pollen DNA metabarcoding using constructed species mixtures. Molecular Ecology 28:431–455.

Bell, K. L., K. S. Burgess, K. C. Okamoto, R. Aranda, and B. J. Brosi. 2016a. Review and future prospects for DNA barcoding methods in forensic palynology. Forensic Science International: Genetics 21:110–116.

Bell, K. L., N. De Vere, A. Keller, R. T. Richardson, A. Gous, K. S. Burgess, and B. J. Brosi. 2016b. Pollen DNA barcoding: current applications and future prospects. Genome 59:629–640.

Bell, K. L., V. M. Loeffler, and B. J. Brosi. 2017. An rbcL reference library to aid in the identification of plant species mixtures by DNA metabarcoding. Appl. Plant Sci. 5:1600110.

Bergstra, J., and Y. Bengio. 2012. Random search for hyper-parameter optimization. The journal of machine learning research 13:281–305.

Bolyen, E., J. R. Rideout, M. R. Dillon, N. A. Bokulich, C. C. Abnet, G. A. Al-Ghalith, H. Alexander, E. J. Alm, M. Arumugam, F. Asnicar, et al. 2019. Reproducible, interactive, scalable and extensible microbiome data science using QIIME 2. Nat. Biotechnol. 37:852–857.

Branco, P., L. Torgo, and R. P. Ribeiro. 2016. A survey of predictive modeling on imbalanced domains. ACM computing surveys (CSUR) 49:1–50.

Breiman, L. 2001. Random forests. Machine learning 45:5–32.

Bryant, V. M., and G. D. Jones. 2006. Forensic palynology: Current status of a rarely used technique in the United States of America. Forensic Science International 163:183–197.

Bryant Jr, V. M., and R. G. Holloway, 1983. The role of palynology in archaeology. Pages 191–224 in Advances in archaeological method and theory. Elsevier.

Callahan, B. J., P. J. McMurdie, M. J. Rosen, A. W. Han, A. J. A. Johnson, and S. P. Holmes. 2016. DADA2: high-resolution sample inference from Illumina amplicon data. Nat. Methods 13:581–583.

Cappellari, A., G. Bonaldi, M. Mei, D. Paniccia, P. Cerretti, and L. Marini. 2022. Functional traits of plants and pollinators explain resource overlap between honeybees and wild pollinators. Oecologia 198:1019–1029.

Chen, T., T. He, M. Benesty, V. Khotilovich, Y. Tang, H. Cho, K. Chen, R. Mitchell, I. Cano, T. Zhou, et al. 2015. Xgboost: extreme gradient boosting. R package version 0.4-2 1:1–4.

Cohen, H., G. P. Smith, H. Sardiñas, J. F. Zorn, Q. S. McFrederick, S. H. Woodard, and L. C. Ponisio. 2021. Mass-flowering monoculture attracts bees, amplifying parasite prevalence. Proceedings of the Royal Society B 288:20211369.

Davis, M. B. 1969. Palynology and environmental history during the Quaternary period. American Scientist 57:317–332.

Di-Giovanni, F., and P. Kevan. 1991. Factors affecting pollen dynamics and its importance to pollen contamination: a review. Canadian Journal of Forest Research 21:1155–1170.

Dietterich, T. G., 2000. Ensemble methods in machine learning. Pages 1–15 in International workshop on multiple classifier systems. Springer.

Du, P., X. Bai, K. Tan, Z. Xue, A. Samat, J. Xia, E. Li, H. Su, and W. Liu. 2020. Advances of four machine learning methods for spatial data handling: A review. Journal of Geovisualization and Spatial Analysis 4:13.

Elger, K., B. K. Biskaborn, H. Pampel, and H. Lantuit. 2016. Open research data, data portals and data publication–an introduction to the data curation landscape. Polarforschung 85:119–133.

Erdtman, G. 2023. Pollen morphology and plant taxonomy: Angiosperms (an introduction to palynology). Brill.

Faegri, K., and J. Iversen. 1992. Textbook of Pollen Analysis. 4 edition. Blackwell Scientific Publications.

Fan Brown, J. J., K. Moriarty, R. McDonald, L. R. Best, J. F. Zorn, and L. C. Ponisio. 2025. Mid-Seral Thinned Stands in Pacific Northwest Coastal Forests Show Enhanced Floral and Bee Diversity without Increased Parasite Prevalence in Wild Bees. J. For. pages 1–27.

Goodman, F., J. Doughty, C. Gary, C. Christou, B. Hu, E. Hultman, D. Deanto, and D. Masters, 2015. PIGLT: a pollen identification and geolocation system for forensic applications. Pages 1–7 in 2015 IEEE International Symposium on Technologies for Homeland Security (HST). IEEE.

Heinrich, B. 1976. The foraging specializations of individual bumblebees. Ecological monographs 46:105–128.

Helderop, E., E. J. Bienenstock, T. H. Grubesic, J. Miller, D. Tong, B. Brosi, and S. Jha. 2021. Network-based geoforensics: Connecting pollen and plants to place. Ecological Informatics 66:101443.

Huang, L., C. Xu, W. Yang, and R. Yu. 2020. A machine learning framework to determine geolocations from metagenomic profiling. Biology direct 15:1–12.

Hwang, G. M., K. C. Riley, C. T. Christou, G. M. Jacyna, J. P. Woodard, R. M. Ryan, M. B. Bush, B. G. Valencia, C. N. McMichael, S. W. Punyasena, et al., 2013. Automated pollen identification system for forensic geo-historical location applications. Pages 297–303 in 2013 IEEE International Conference on Technologies for Homeland Security (HST). IEEE.

Karstens, L., M. Asquith, S. Davin, D. Fair, W. T. Gregory, A. J. Wolfe, J. Braun, and S. McWeeney. 2019. Controlling for contaminants in low-biomass 16S rRNA gene sequencing experiments. MSystems 4.

Katoh, K., and D. M. Standley. 2013. MAFFT multiple sequence alignment software version 7: improvements in performance and usability. Mol. Biol. Evol. 30:772–780.

Keller, A., N. Danner, G. Grimmer, v. d. Ankenbrand, M, K. Von Der Ohe, W. Von Der Ohe, S. Rost, S. Härtel, and I. Steffan-Dewenter. 2015. Evaluating multiplexed next-generation sequencing as a method in palynology for mixed pollen samples. Plant Biology 17:558–566.

Kembel, S. W., T. K. O’Connor, H. K. Arnold, S. P. Hubbell, S. J. Wright, and J. L. Green. 2014. Relationships between phyllosphere bacterial communities and plant functional traits in a neotropical forest. Proc. Natl. Acad. Sci. U.S.A 111:13715–13720.

Lee, S. J., H. Liu, and M. D. Ward. 2019. Lost in space: Geolocation in event data. Political science research and methods 7:871–888.

Lemanski, N. J., C. N. Cook, B. H. Smith, and N. Pinter-Wollman. 2019. A multiscale review of behavioral variation in collective foraging behavior in honey bees. Insects 10:370.

McFrederick, Q. S., and S. M. Rehan. 2016. Characterization of pollen and bacterial community composition in brood provisions of a small carpenter bee. Molecular ecology 25:2302–2311.

McGlone, M. S., 1988. New Zealand. Pages 557–599 in B. Huntley and T. I. Webb, editors. Vegetation History. Kluwer Academic Publishers, Amsterdam.

Mildenhall, D., P. E. Wiltshire, and V. M. Bryant. 2006. Forensic palynology: why do it and how it works. Forensic science international 163:163–172.

Milusheva, S., R. Marty, G. Bedoya, S. Williams, E. Resor, and A. Legovini. 2021. Applying machine learning and geolocation techniques to social media data (Twitter) to develop a resource for urban planning. PloS one 16:e0244317.

Morris, W. F. 1993. Predicting the consequence of plant spacing and biased movement for pollen dispersal by honey bees. Ecology 74:493–500.

Omonhinmin, C., and C. Onuselogu. 2022. rbcL gene in global molecular data repository. Data in Brief 42:108090.

Pedregosa, F., G. Varoquaux, A. Gramfort, V. Michel, B. Thirion, O. Grisel, M. Blondel, P. Prettenhofer, R. Weiss, V. Dubourg, J. Vanderplas, A. Passos, D. Cournapeau, M. Brucher, M. Perrot, and E. Duchesnay. 2011. Scikit-learn: Machine Learning in Python. Journal of Machine Learning Research 12:2825–2830.

Price, M. N., P. S. Dehal, and A. P. Arkin. 2010. FastTree 2–approximately maximum-likelihood trees for large alignments. PloS One 5:e9490.

Qi, Y. 2012. Random forest for bioinformatics. Ensemble machine learning: Methods and applications pages 307–323.

Ranstam, J., and J. A. Cook. 2018. LASSO regression. Journal of British Surgery 105:1348–1348.

Rimal, Y., N. Sharma, and A. Alsadoon. 2024. The accuracy of machine learning models relies on hyperparameter tuning: student result classification using random forest, randomized search, grid search, bayesian, genetic, and optuna algorithms. Multimedia Tools and Applications 83:74349–74364.

Smith, G. P., H. Cohen, J. F. Zorn, Q. S. McFrederick, and L. C. Ponisio. 2024. Plant– pollinator network architecture does not impact intraspecific microbiome variability. Molecular Ecology 33:e17306.

Smith, G. P., J. Gardner, J. Gibbs, T. Griswold, M. Hauser, D. Yanega, and L. C. Ponisio. 2021. Sex-associated differences in the network roles of pollinators. Ecosphere 12:e03863.

Stajich, J. E., D. Block, K. Boulez, S. E. Brenner, S. A. Chervitz, C. Dagdigian, G. Fuellen, J. G. Gilbert, I. Korf, H. Lapp, et al. 2002. The Bioperl toolkit: Perl modules for the life sciences. Genome Res. 12:1611–1618.

Steinbach, M., and P.-N. Tan, 2009. kNN: k-nearest neighbors. Pages 165–176 in The top ten algorithms in data mining. Chapman and Hall/CRC.

Testas, A., 2023. Decision tree regression with pandas, scikit-learn, and PySpark. Pages 75–113 in Distributed Machine Learning with PySpark: Migrating Effortlessly from Pandas and Scikit-Learn. Springer.

Thessen, A. 2016. Adoption of machine learning techniques in ecology and earth science. One Ecosystem 1:e8621.

Tong, D., T. H. Grubesic, W. Mu, J. A. Miller, E. Helderop, S. Jha, B. J. Brosi, and E. J. Bienenstock. 2021. Identifying the spatial footprint of pollen distributions using the Geoforensic Interdiction (GOFIND) model. Computers, Environment and Urban Systems 87:101615.

Visick, O. D., and F. L. Ratnieks. 2023. Density of wild honey bee, Apis mellifera, colonies worldwide. Ecology and Evolution 13:e10609.

Warny, S., S. Ferguson, M. S. Hafner, and G. Escarguel. 2020. Using museum pelt collections to generate pollen prints from high-risk regions: a new palynological forensic strategy for geolocation. Forensic science international 306:110061.

Weber, M., and S. Ulrich. 2017. PalDat 3.0–second revision of the database, including a free online publication tool. Grana 56:257–262.

Williams, J. W., and B. Shuman. 2008. Obtaining accurate and precise environmental reconstructions from the modern analog technique and North American surface pollen dataset. Quaternary Science Reviews 27:669–687.

Wiltshire, P. E. 2016. Protocols for forensic palynology. Palynology 40:4–24.

Yang, Y., B. Xie, and J. Yan. 2014. Application of next-generation sequencing technology in forensic science. Genomics, proteomics & bioinformatics 12:190–197.

Young, A. M., P. L. Kohl, B. Rutschmann, I. Steffan-Dewenter, A. Brockmann, and F. C. Dyer. 2021. Temporal and spatial foraging patterns of three Asian honey bee species in Bangalore, India. Apidologie 52:503–523.

Zhang, C., and Y. Ma. 2012. Ensemble machine learning. Springer.

