## SupplementaryInformation for "Assessing the potential of bee-collected pollen sequence data to train machine learning models for geolocation of sample origin"

### Supplementary Information

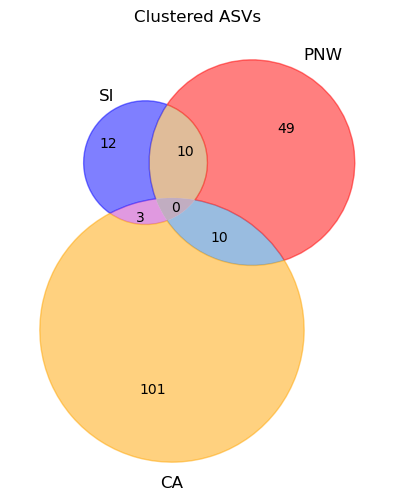

Figure S1: The overlap between projects of taxonomically clustered ASVs. Projects are encoded as follows: California Sunflowers Project (CA), Pacific Northwest Forests Project (PNW), and Sky Islands Project (SI).

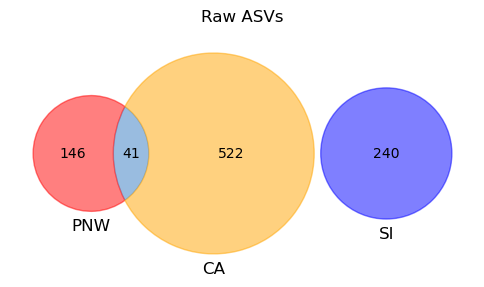

Figure S2: The overlap between projects of raw ASVs. Projects are encoded as follows: California Sunflowers Project (CA), Pacific Northwest Forests Project (PNW), and Sky Islands Project (SI).

Table S1: Hyperparameter search space for each model.

| **Model** | **Hyperparameter Search Space** |
| --- | --- |
| MultiTaskLasso | alpha: [0.00, 1.0) step 0.01 max_iter = 10,000 |
| SVR | C: Uniform(0.1, 10) kernel: [linear, rbf, poly] gamma: [scale, auto] |
| KNN | n_neighbors: 2–49 weights: [uniform, distance] p: [1, 2] algorithm: [auto] |
| Decision Tree | max_depth: None + [3–20] min_samples_split: [2, 5, 10, 20, 50] min_samples_leaf: [1, 2, 5, 10, 20, 50] max_features: [auto, sqrt, log2, None] max_leaf_nodes: None + [10–100, step 10] |
| Random Forest | n_estimators: [50, 100, 200, 500, 1000] max_depth: None + [10, 20, 30, 40, 50] min_samples_split: [2, 5, 10, 20, 50] min_samples_leaf: [1, 2, 5, 10, 20, 50] max_features: [auto, sqrt, log2, None] |
| XGBoost | n_estimators: [100, 200, 500, 1000] learning_rate: [0.01, 0.05, 0.1, 0.2, 0.3] max_depth: [3, 5, 7, 10, 15, 20] min_child_weight: [1, 5, 10, 20] subsample: [0.6, 0.7, 0.8, 0.9, 1.0] colsample_bytree: [0.6, 0.7, 0.8, 0.9, 1.0] gamma: [0, 0.1, 0.2, 0.5, 1, 5] reg_alpha: [0, 0.01, 0.1, 1, 10, 100] reg_lambda: [1, 10, 50, 100] |

Table S2: Best-tuned hyperparameters for taxonomic and raw models

| **Model** | **Taxonomic Model Features** | **Raw Model Features** |
| --- | --- | --- |
| MultiTaskLasso | alpha=0.08 | alpha=0.08 |
| SVR | C=2.847, gamma=auto, kernel=rbf | C=2.847, gamma=auto, kernel=rbf |
| KNN | weights=uniform, p=2, n_neighbors=3, algorithm=auto | weights=uniform, p=2, n_neighbors=3, algorithm=auto |
| Decision Tree | min_samples_split=10, min_samples_leaf=2, max_leaf_nodes=40, | min_samples_split=50, min_samples_leaf=1, max_leaf_nodes=None, |
| Random Forest | max_features=None, max_depth=18, random_state=seed | max_features=sqrt, max_depth=None, random_state=seed |
| XGBoost |  |  |
|  | n_estimators=200, min_samples_split=5, min_samples_leaf=1, max_features=log2, max_depth=None | n_estimators=200, min_samples_split=5, min_samples_leaf=1, max_features=log2, max_depth=None |
|  | subsample=0.8, reg_lambda=1, reg_alpha=1, n_estimators=500, | subsample=0.8, reg_lambda=1, reg_alpha=1, n_estimators=500, |
|  | min_child_weight=1, max_depth=20, learning_rate=0.3, gamma=0, colsample_bytree=0.8 | min_child_weight=1, max_depth=20, learning_rate=0.3, gamma=0, colsample_bytree=0.8 |

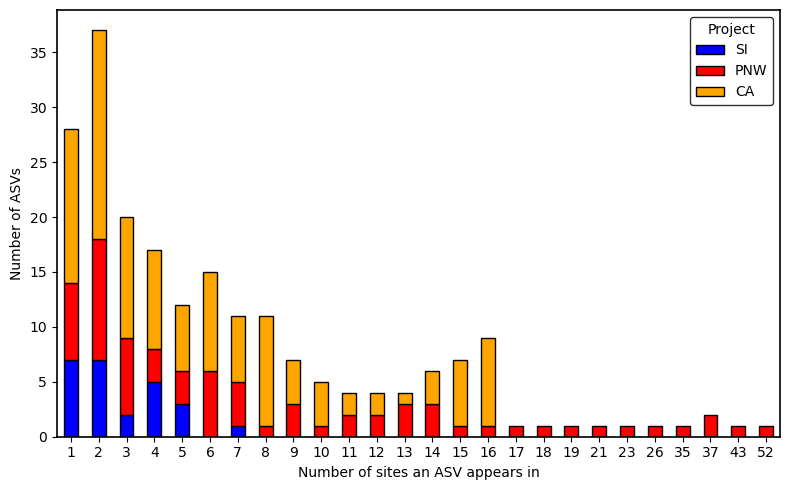

Figure S3: Distribution of taxonomically clustered ASV occurrence across sites by project. An ASV was considered present at a site if it was detected in at least one individual at a given site for that project. Projects are encoded as follows: California Sunflowers Project (CA), Pacific Northwest Forests Project (PNW), and Sky Islands Project (SI).

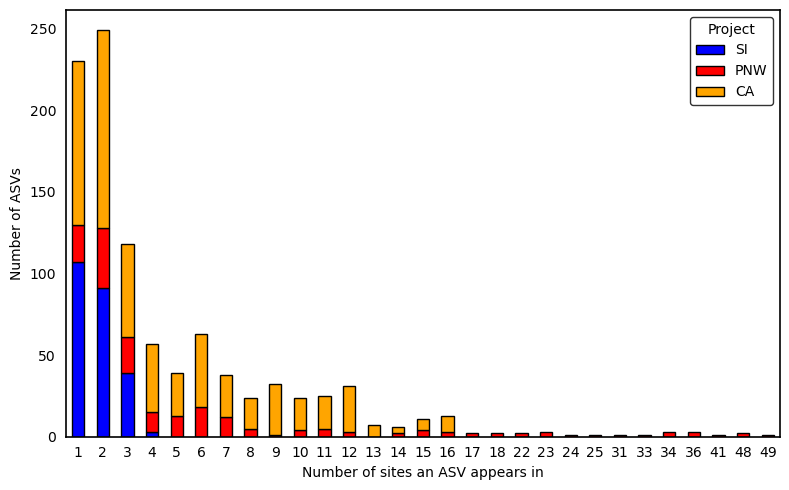

Figure S4: Distribution of raw ASV occurrence across sites by project. An ASV was considered present at a site if it was detected in at least one individual at a given site for that project. Projects are encoded as follows: California Sunflowers Project (CA), Pacific Northwest Forests Project (PNW), and Sky Islands Project (SI).

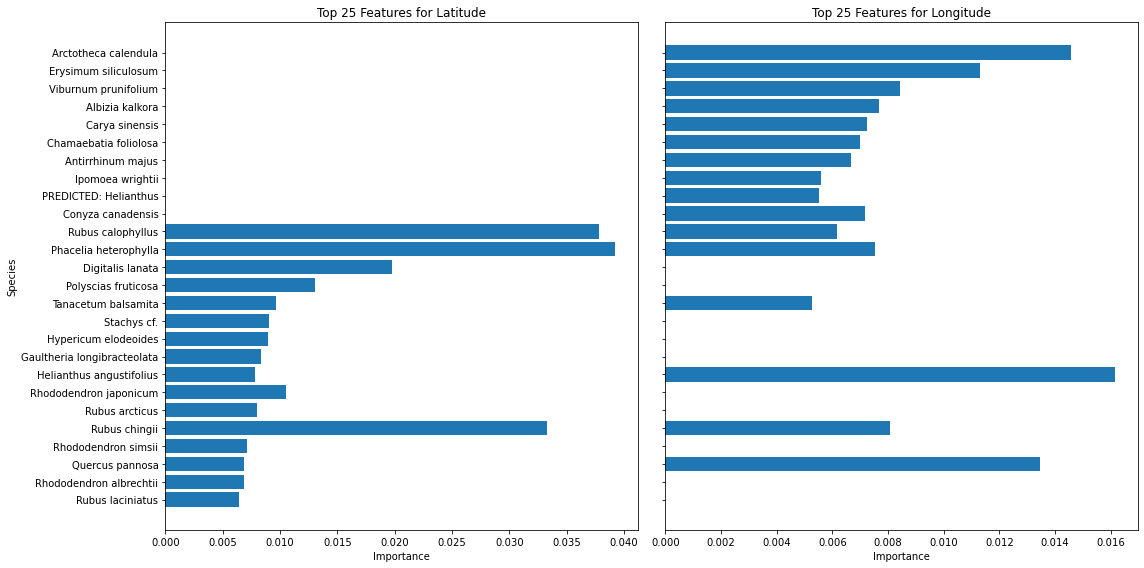

*Figure S5: Feature importances for top 25 most informative ASVs for the models trained on raw sequence data. Feature importances were extracted and then classified taxonomically using NCBI BLAST.*

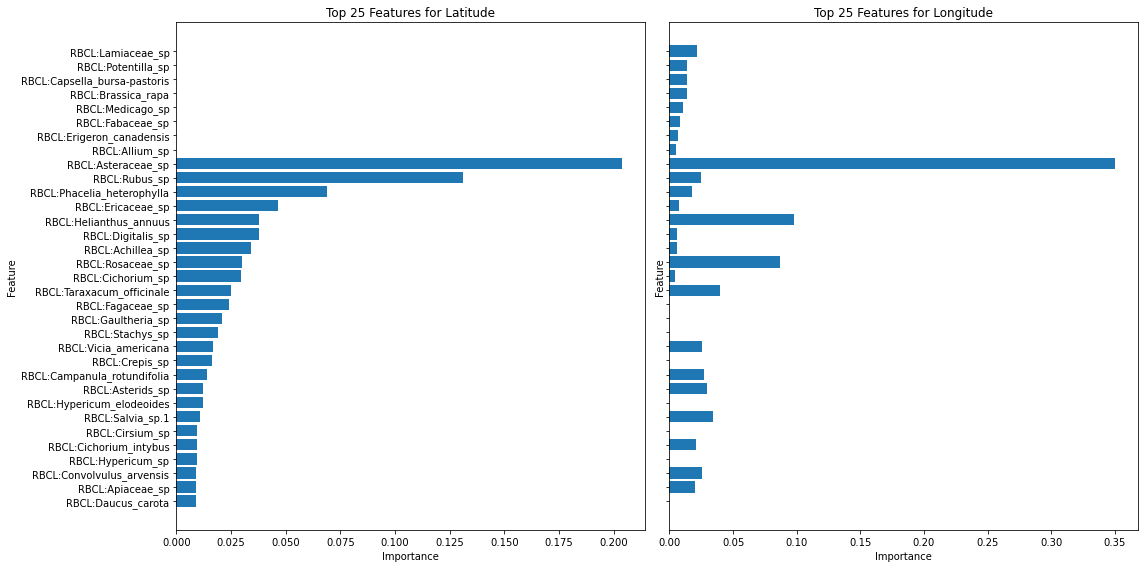

*Figure S6: Feature importances for top 25 most informative ASVs for the models trained on taxonomically-assigned data.*
